# Phylogeny and Species Delimitation in *Isoxylosteum*, a *Lonicera* Clade Endemic to the Himalayan-Tibetan-Hengduan Region

**DOI:** 10.64898/2026.08.31.748305

**Authors:** Mansa Srivastav, Amit Kumar, Gopal Singh Rawat, Richard H. Ree, Michael J. Donoghue

## Abstract

The Himalayan-Tibetan-Hengduan (HTH) region is the richest biodiversity hotspot for high-elevation plants. However, owing to its remote and physically challenging topography as well as the trans-national nature of the region, many taxonomic problems in the area remain unresolved, particularly in the Himalaya. This, in turn, has impeded our understanding of the assembly of its extraordinary high-elevation flora. Here, we resolve phylogenetic relationships and delimit species in a distinctive clade of honeysuckles that is endemic to the HTH – the *Isoxylosteum* clade of *Lonicera –* using restriction-site associated DNA sequencing (RADseq) and morphological data. Five species complexes of *Isoxylosteum* have standardly been recognized. Three of these complexes are highly variable and have been divided into several varieties or species each. Phylogenetic, population structure, and morphological analyses of leaf and floral traits from samples collected across the range of the clade support the recognition of five species, including a species that has most often been recognized as a variety of *L. rupicola* (*L. rupicola* var. *minuta*). Instead, we find that it is sister to *L. spinosa*. This is surprising because the geographic range of *L. minuta* is contiguous with the other varieties of *L. rupicola* in the northern Hengduan region but widely separated from *L. spinosa* whose range lies mainly to the west of the Tibetan plateau. On close examination we find that several morphological and ecological traits also support a closer relation of *L. minuta* to *L. spinosa*. None of the other eight previously recognized varieties and species were supported. Floral traits showed high discriminatory power, correctly classifying 89% of samples to species. By comparison, leaf dimensions classified species with 59% accuracy. Our results identify diagnostic morphological apomorphies for each recognized species and major clade and provide a revised taxonomic framework for *Isoxylosteum*.

---

The Himalayan-Tibetan-Hengduan (HTH) mountain-plateau system began forming ∼50-55 million years ago following the collision of the Indian and Eurasian plates, creating the highest mountains in the world (Favre et al. 2015). Today, it harbors the world’s richest temperate alpine flora (Ding et al. 2020). Many high-elevation plant lineages are endemic to the HTH (Zhang et al. 2016; Wambulwa 2021), and could provide insights into how its extraordinary flora was assembled. However, due to the difficulty of accessing its remote areas, coupled with the fact that this vast system straddles multiple political boundaries (Fig. 1), studies sampling lineages across their entire geographic ranges remain rare, especially those sampling Himalayan populations (Wambulwa 2021). Consequently, many taxonomic problems remain unresolved in the HTH. This has hindered our understanding of the origin of its high-altitude flora, the pathways and barriers underlying the dispersal of these lineages, and the evolutionary basis of their adaptations to the diverse high-altitude biomes encompassed by the HTH.

**Figure 1.**
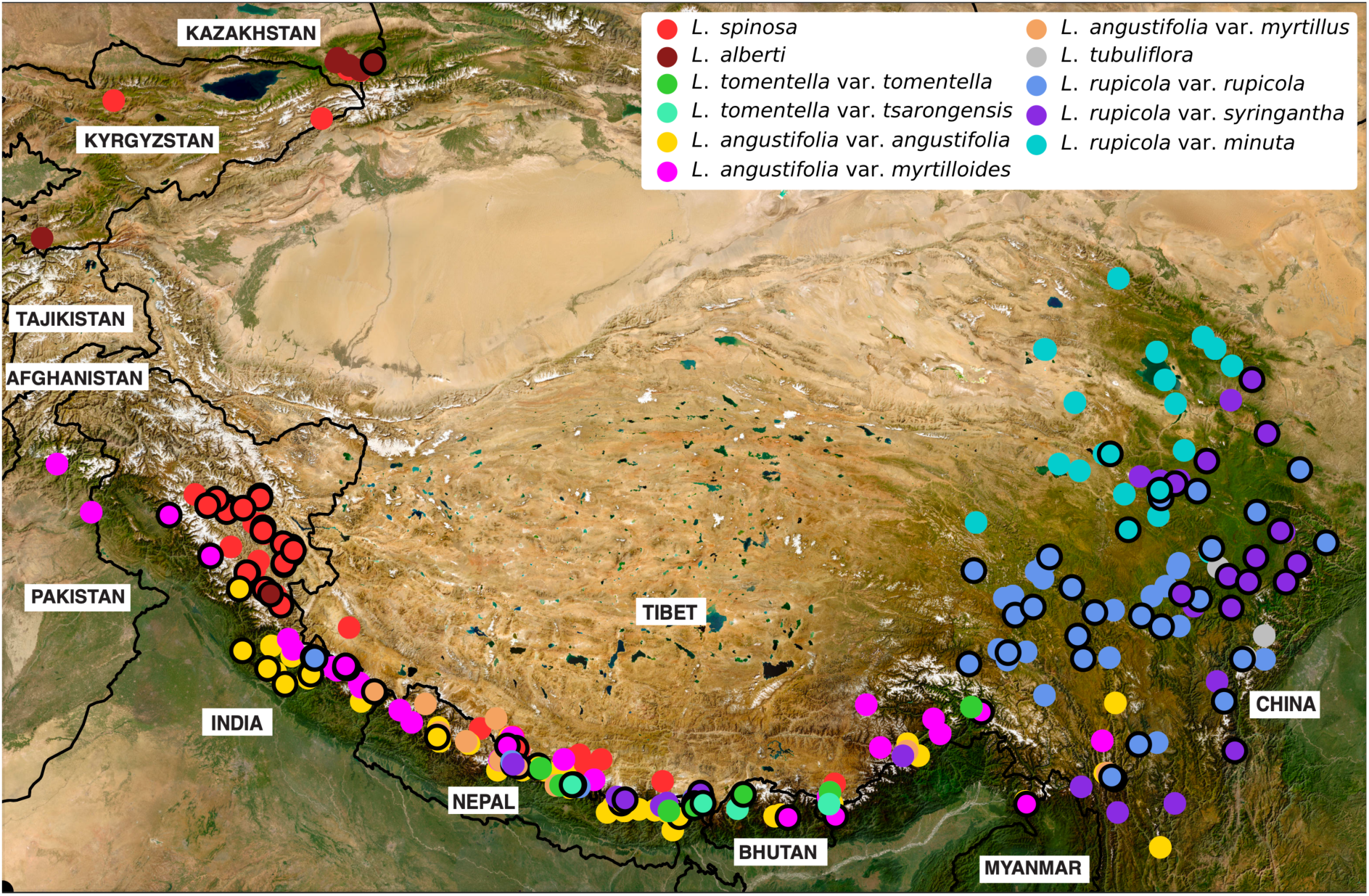
Map of the Himalayan-Tibetan-Hengduan region showing the distribution of the currently recognized species and varieties of *Isoxylosteum*. The country labels illustrate the trans-boundary nature of this mountain-plateau system. Populations sampled in this study are circled in black.

Here, we address this problem by resolving the phylogeny and taxonomy of the *Isoxylosteum* clade, a small but highly variable HTH-endemic group of honeysuckles (*Lonicera* L., Caprifoliaceae, Dipsacales) that spans the geographic regions and biomes of this mountain–plateau system. *Isoxylosteum* was formally recognized as a section within *Lonicera* by Rehder (1903) and has been strongly supported as a clade in a series of phylogenetic analyses (Theis et al. 2008; Smith 2009; Nakaji et al. 2015; Srivastav et al. 2023). *Isoxylosteum* species are unique within *Lonicera* in having radially symmetrical flowers with five nectaries, and leaves that are folded at the midrib in bud, as opposed to the ancestral condition of having bilaterally symmetrical flowers with one to three nectaries and leaves with convolute or involute vernation (Rehder 1903; Pusalkar 2011; Srivastav et al. 2023). For these reasons, it was elevated to genus rank as *Devendraea* by Pusalkar (2011). However, as *Isoxylosteum* is nested well within *Lonicera*, its recognition as a separate genus would render *Lonicera* paraphyletic. Specifically, *Isoxylosteum* is placed within the large subgenus *Chamaecerasus*, sister to a clade containing the whole of *Lonicera* section *Coeloxylosteum* plus the *Rhodantheae* subsection of the paraphyletic section *Isika* (Srivastav et al. 2023).

There have been two range-wide taxonomic treatments of *Isoxylosteum.* These have recognized five to seven species, and several varieties (Pusalkar et al. 2011; Yang et al. 2011). Some taxonomic entities recognized as species in one treatment have been treated as varieties in others, or not recognized at all. In general, however, five major species complexes have been recognized within *Isoxylosteum,* as follows:

1. *Lonicera spinosa* (Decne.) Jacquem. ex Walp. occupies the cold deserts of western and central Himalaya as well as those of the Pamiro-Alai and Tien Shan mountains of Central Asia (Fig. 1). *L. alberti* Regel, described from Central Asia, has been differentiated from *L. spinosa* by its lack of thorns and ovate-elliptic corolla lobes, in contrast to the thorny habit and lanceolate-obtuse corolla lobes of *L. spinosa*. Pusalkar (2011) recognized *L. alberti* as a separate species while Yang et al. (2011) synonymized it with *L. spinosa*.
2. *Lonicera tomentella* Hook.f. & Thomson occurs in temperate to subalpine habitats from Nepal in central Himalaya to eastern Himalaya as well as adjacent regions of Tibet. Two varieties have been recognized within this complex based on differences in leaf traits: *L. tomentella* var. *tomentella* and *L. tomentella* var. *tsarongensis*. While var. *tomentella* has leaves that are non-glaucous and pubescent abaxially, with ∼7-8 pairs of lateral veins, var. *tsarongensis* leaves are less hairy and can be glaucous, with ∼4-5 pairs of lateral veins (Pusalkar 2011; Yang et al. 2011).
3. *Lonicera rupicola* Hook. f. & Thomson is a highly variable complex. Yang et al. (2011) recognized three varieties within it. In *L. rupicola* var. *rupicola* the leaves are covered abaxially with dense, white hair, whereas *L. rupicola* var. *syringantha* has glabrous leaves. Both varieties are found in the Hengduan and the Himalaya in alpine areas, cold deserts, and forest margins. The third variety, *L. rupicola* var. *minuta*, differs from the other two in habitat—it is found in shifting sands in the cold deserts of Qinghai and Gansu (Batalin 1892; Rehder 1903; Yang et al. 2011). Pusalkar (2011) treated *L. minuta* as a species within section *Rupicolae* along with *L. rupicola* var. *rupicola* and *L. rupicola* var. *syringantha*.
4. *Lonicera angustifolia* Wall. ex DC. is also a highly variable complex. Yang et al. (2011) recognize two varieties that differ in leaf size and shape, and peduncle length: *L. angustifolia* var. *angustifolia* has larger leaves, with acute apices, and long peduncles while *L. angustifolia* var. *myrtillus* (J. D. Hooker & Thomson) Q. E. Yang, Landrein, Borosova & J. Osborne has small leaves, with obtuse apices and short peduncles. Pusalkar (2011) treated these two varieties as separate species and additionally recognized four varieties within *L. myrtillus* on the basis of leaf and bract shape differences (var. *cyclophylla,* var*. depressa,* var. *minutifolia*, and var. *myrtillus*). A third variety has been described within the *L. angustifolia* complex, *L. angustifolia* var. *myrtilloides* Purpus. It has leaf and peduncle lengths intermediate between *L. angustifolia* var. *angustifolia* and *L. angustifolia* var. *myrtillus* and leaf apices that vary from acute to obtuse (Rehder 1903; Pusalkar 2011; Yang et al. 2011).
5. *Lonicera tubuliflora* Rehder is a narrow endemic reported only from a few mid-elevation localities in Sichuan, China. It closely resembles *L. angustifolia* but differs in having free ovaries and narrow tubular corollas with lobes ∼1.5 mm (Yang et al. 2011). This species was not treated by Pusalkar (2011).

Previous studies of *Lonicera* phylogeny sampled just a single individual per species/variety from four of the five species complexes (Theis et al. 2008; Smith 2009; Srivastav et al. 2023). These individuals were found in all analyses to form a well-supported clade, within which *L. spinosa* plus *L. rupicola* formed a clade sister to *L. tomentella* plus *L. angustifolia.* Here, we sampled multiple individuals of four of the five species complexes and all of the recognized varieties within them from localities across their geographic ranges (except the four varieties of *L. myrtillus* recognized only by Pusalkar 2011, and synonymized elsewhere, Yang et al. 2011). For the rare *L. tubuliflora,* we were able to include only one specimen.

We combine restriction-site associated DNA sequencing (RADseq) with analyses of a suite of morphological traits to resolve phylogenetic relationships, delimit independently evolving species, and identify diagnostic morphological apomorphies for the recognized species and major clades. Together, these analyses show that *Isoxylosteum* is a small, well-supported, and highly distinctive lineage that has spread throughout the Himalaya and Hengduan mountains and around the Tibetan Plateau. We found support for the recognition of five species, including one previously treated as a variety of *L. rupicola* that is instead recovered as sister to the geographically distant *L. spinosa*. None of the other varieties and species were found to form independently evolving units. Our findings provide a robust systematic framework for *Isoxylosteum* and establish it as a model lineage for investigating the assembly and adaptation of the high-altitude flora of the HTH. They also provide a phylogenetic framework for future studies of floral evolution and pollination biology. In a companion paper in preparation, we leverage this information to investigate the past and present climatic and geographic factors underlying the origin and maintenance of the geographic ranges of species established here, including the barriers that restrict movement and gene flow among species.

## Materials and methods

### Sampling and DNA Extraction

We sampled 103 individuals belonging to the five previously recognized species from throughout their ranges (Fig.1). Samples were collected in the field from India and Nepal from 2018 to 2022, as well as from herbarium specimens and silica gel material from the Biodiversity of the Hengduan mountains project (http://hengduan.huh.harvard.edu/fieldnotes) and several herbaria (see Appendix 1). Our sample includes 18 accessions assigned to *L. spinosa*, two to *L. alberti*, 45 to *L. rupicola*, 31 to *L. angustifolia*, six to *L. tomentella*, and one to *L. tubuliflora*. For the three large species complexes, *L. rupicola*, *L. angustifolia*, and *L. tomentella*, we sampled all varieties recognized within each—*L. rupicola* var. *rupicola* (28 accessions), *L. rupicola* var. *syringantha* (14), and *L. rupicola* var. *minuta* (3); *L. angustifolia* var. *angustifolia* (15), *L. angustifolia* var. *myrtillus* (14), and *L. angustifolia* var. *myrtilloides* (2); *L. tomentella* var. *tomentella* (3), and *L. tomentella* var. *tsarongensis* (3). Our samples largely span the geographic range of the clade, though we have only one sample from Central Asia where *L. alberti* and *L. spinosa* occur. We were able to obtain only one individual of the rare *L. tubuliflora*; we note that this was from Sikkim, Eastern Himalaya, outside of its previously reported range in Sichuan as COVID-19 restrictions prevented field work and acquisition of material from China. We included two outgroup species, *L. arborea* and *L. oblongifolia* from sections *Coeloxylosteum* and *Isika*, respectively (Srivastav et al. 2023; Srivastav and Donoghue, MS in prep.). Genomic DNAs were extracted using the Beck et al. (2012) modification of the standard CTAB protocol (Doyle and Doyle 1987) and cleaned using SpeedBeads (Rohland and Reich 2012).

### Library preparation and data assembly

RADseq libraries were prepared by Floragenex, Beaverton, Oregon, USA (http://floragenex.com). Samples were digested using the PstI enzyme followed by adaptor ligation, sonication, end-repair, purification, PCR amplification, and size selection for fragments of length 400–600 base pairs (bp). The average fragment length was 496 bp. Our samples were distributed over three plates. The first two plates were sequenced on an Illumina HiSeq4000, while the third plate was sequenced on a NovaSeq6000, all at the University of Oregon GC3F facility (http://gc3f.uoregon.edu).

Loci were assembled using ipyrad version 0.9.92 (Eaton and Overcast 2020), using the whole-genome assembly of *Lonicera japonica* as a reference (Pu et al. 2020). To infer the optimal clustering threshold, we evaluated a range of thresholds from 85% to 96% and identified the value that maximised the number of parsimony-informative sites (PIS), mean number of loci, total base pairs, and SNPs. A threshold of 0.91 produced the highest values for all metrics except the number of SNPs and was therefore used for all subsequent analyses. To test the impact of missing data, we created four assemblies: i) min4 that retained l66,463 loci across at least four samples with 40.92% missing data, ii) min10 that retained 55,863 loci across at least 10 samples with 38.83% missing data, iii) min20 that retained 49,462 loci across at least 20 samples with 36.91% missing data, and iv) min40 that retained 37,264 loci across at least 40 samples with 32.10% missing data.

### Phylogenomic analyses

We inferred trees from the concatenated datasets using IQ-TREE version 2.2.2.7 (Minh et al. 2020) with 1000 ultrafast bootstrap replicates (Hoang et al. 2018) under the K3Pu+F+I+R10 model. The model was selected using ModelFinder (Kalyaanamoorthy et al. 2017) based on Bayesian Information Criterion (BIC). Quartet-based trees were reconstructed using Tetrad version 0.9.13 in ipyrad (Eaton and Overcast 2020). We sampled all possible quartets and ran 100 bootstrap replicates. Trees were visualized using Figtree v. 1.4.4 (Rambaut 2012).

### Population Genetic Analyses

#### Structure analyses

We used the Bayesian clustering method STRUCTURE to identify genetic clusters within *Isoxylosteum* (Pritchard et al. 2000), using the ipyrad-analysis toolkit (Eaton and Overcast 2020). We retained loci shared across at least 85% of the samples, which yielded 2,497 unlinked SNPs. Analyses were run at K values ranging from 1-11, covering all previously recognized species and varieties. Ten replicates were run for 1,000,000 generations with a burn-in of 100,000. Outputs were permuted using CLUMPP v1.1.2 (Jakobsson and Rosenberg 2007) in ipyrad. The optimal K value was extracted and plotted in ipyrad, and estimated using ΔK as recommended by Evanno et al. (2015). Since STRUCTURE is prone to identifying only the highest-order pattern (Jombart et al. 2010), we subsequently ran analyses on each of the identified clusters, retaining loci shared across at least 70% of the samples.

#### Discriminant Analysis of Principal Components

We inferred population structure using the multivariate method, discriminant analysis of principal components (DAPC), in the R package *adegenet* (Jombart 2008). DAPC uses Principal Component Analysis (PCA) followed by discriminant analysis to identify clusters (Jombart et al. 2010). We used the same assemblies as in the STRUCTURE analyses. The number of clusters was identified using the function *find.clusters*. ‘Elbow’ of the BIC plot was selected as the optimal *K* value and *a*-score was used for determining the number of principal components to retain.

#### Principal Component Analysis of snps

We analyzed SNPs using PCA with the min4 assembly in the ipyrad toolkit. Missing data were imputed using two approaches: (1) a sample-based method in which samples were assigned to predefined groups based on phylogenetic results; and (2) a k-means clustering method that inferred groups without *a priori* assignments. For both approaches, we used a minimum coverage threshold (mincov) of 0.75 and a minimum mapping threshold (minmap) of 0.50 to further retain SNPs with data for at least 50% of the samples within each group, and conducted 25 replicates. For method (1) we assigned samples to the five clades inferred by phylogenetic analyses, resulting in 15,612 unlinked SNPs; for method (2) we used a K value of 5 that resulted in 1,564 unlinked SNPs.

#### Pairwise FST

To estimate genetic differentiation between populations, we calculated pairwise F_ST_ statistics between the five recovered clades (Holsinger and Weir 2009) using VCFTools v0.1.6 (Danecek et al. 2011) based on the Weir and Cockerham estimator (Weir and Cockerham 1984).

#### Bpp analyses

We performed species delimitation in BPP v.4.6.2 using the A10 analysis (Flouri et al. 2018). This analysis uses reversible-jump MCMC to explore species delimitation models compatible with a fixed guide tree (Yang 2015). We subsampled three to seven geographically spaced samples from each of the five main clades recovered in phylogenetic analyses, using 500 loci for the analyses. The guide tree was based on our phylogenetic and STRUCTURE analyses. We used a uniform rooted-tree prior for the species-delimitation models. Analyses were run at two sets of theta (population size parameter) and tau (root age) values: θ ∼ IG(3,0.002), T ∼ IG(3,0.015), and θ ∼ IG(3,0.02), T ∼ IG(3,0.308) that allowed shallower divergence times with smaller population sizes, and deeper divergence times with larger population sizes, respectively. Each analysis was conducted using Algorithm 0 with ε = 2 and Algorithm 1 with α = 2 and m = 1. We ran two replicates, each of length 100,000 generations with a burn-in of 2,000, sampling every 50 generations. Convergence was assessed by verifying that runs with different starting delimitations produced the same results. ESS values were checked in Tracer (Rambaut et al. 2018). We also conducted the A11 analysis, which jointly performs species tree inference and species delimitation, under the same θ and τ prior settings.

### Morphological analyses

We measured leaf and flower traits from herbarium specimens using ImageJ (Schneider et al. 2012). One leaf and one flower were measured per specimen. Floral traits were measured from a total of 146 specimens: 53 of *L. angustifolia*, 36 of *L. rupicola*, 25 of *L. spinosa*, 22 of *L. tomentella*, 6 of *L. tubuliflora*, and 4 of *L. minuta*. For leaf traits, the sampling of *L. tubuliflora* and *L. minuta* was increased to 21 and 7, respectively. Note that we performed analyses with samples identified as *L. tubuliflora* as well, despite the fact that our analyses do not support its recognition as a separate species, but rather its inclusion within *L. angustifolia*. Additional samples are needed to further test these relationships. However, in the meantime, we wanted to explore how its leaf and flower measurements compared to *L. angustifolia* and the other species recognized here.

Leaf length and width were measured from leaves sampled at third or fourth nodes from the tip of a branch, as all species bore fully expanded, mature leaves from that node onwards. We quantified 13 floral traits: peduncle length, corolla length, corolla diameter (the maximum spread of corolla lobes), corolla tube length, corolla tube width at midpoint, calyx tube length, calyx tube width at base, calyx lobe length and width, corolla lobe length and width, and ovary length and breadth. We used PCA to summarize patterns of morphological variation and linear discriminant analysis (LDA) to assess the ability of the traits to distinguish among species, using R (R Core Team 2025). Species differences were assessed using Welch’s one-way ANOVA followed by Games–Howell post hoc tests to account for unequal variances and sample sizes. PCA was performed prior to LDA, and the resulting principal component (PC) scores were used as input variables for the discriminant analysis. LDA was performed using the *lda* function in the MASS package (Ripley and Venables 2002). We used ggplot2 for visualizing results and preparing figures (Wickham 2016).

### Ancestral state reconstructions

We reconstructed the evolution of 11 categorical morphological traits that vary within *Isoxylosteum*. Characters and their states are described in Appendix S1. Reconstructions were based on three geographically spaced accessions of each of the five major clades identified in our phylogenetic and STRUCTURE analyses. Trait data were compiled for the five *Isoxylosteum* clades and 112 other *Lonicera* species (Srivastav and Donoghue, MS in prep.). Reconstructions were performed in a maximum likelihood framework using phytools v1.0-1 (Revell 2012) on a well-resolved IQ-TREE.

### Occurrence and elevation data

We obtained GPS coordinates and elevation data for 383 accessions by compiling information from our own collections; the Biodiversity of the Hengduan mountains project (http://hengduan.huh.harvard.edu/fieldnotes); online biodiversity databases, namely, the Global Biodiversity Information Facility (GBIF; https://www.gbif.org/), iNaturalist (https://www.inaturalist.org/), and the Plant Photo Bank of China (PPBC; http://ppbc.iplant.cn/; Xie et al. 2024); and herbarium specimens deposited at the Botanical Survey of India at Dehradun (BSD), Itanagar (ARUN), and Gangtok (BSI-SHRC), the Herbarium of the University of Kashmir (KASH), the National Herbarium and Plant Laboratories, Kathmandu (KATH), the Moscow State University Herbarium (MW; https://plant.depo.msu.ru/), the New York Botanical Garden Herbarium (NYBG), the Missouri Botanical Garden Herbarium (MO), the Yale Herbarium (YU), the Herbarium of the Royal Botanic Gardens Kew (K), the Royal Botanic Garden Edinburgh Herbarium (E), the John G. Searle Herbarium of the Field Museum of Natural History (F), the Harvard University Herbaria (HUH), herbarium at the Kunming Institute of Botany, Chinese Academy of Sciences (KUN), and the Chinese Virtual Herbarium (CVH; https://www.cvh.ac.cn/).

## Results

### Phylogenomic analyses

Our concatenated and coalescent trees were highly congruent in recovering five major, well-supported clades (bootstrap support ≥ 99) across different assemblies (Fig. 2). As in previous phylogenetic studies of *Lonicera*, *L. spinosa* and *L. rupicola* are recovered as sister to a clade comprising *L. tomentella* and *L. angustifolia*. Surprisingly, in disagreement with all prior taxonomic treatments, *L. rupicola* var. *minuta* is not linked with the other *L. rupicola* varieties but instead is inferred to be sister to *L. spinosa*.

**Figure 2.**
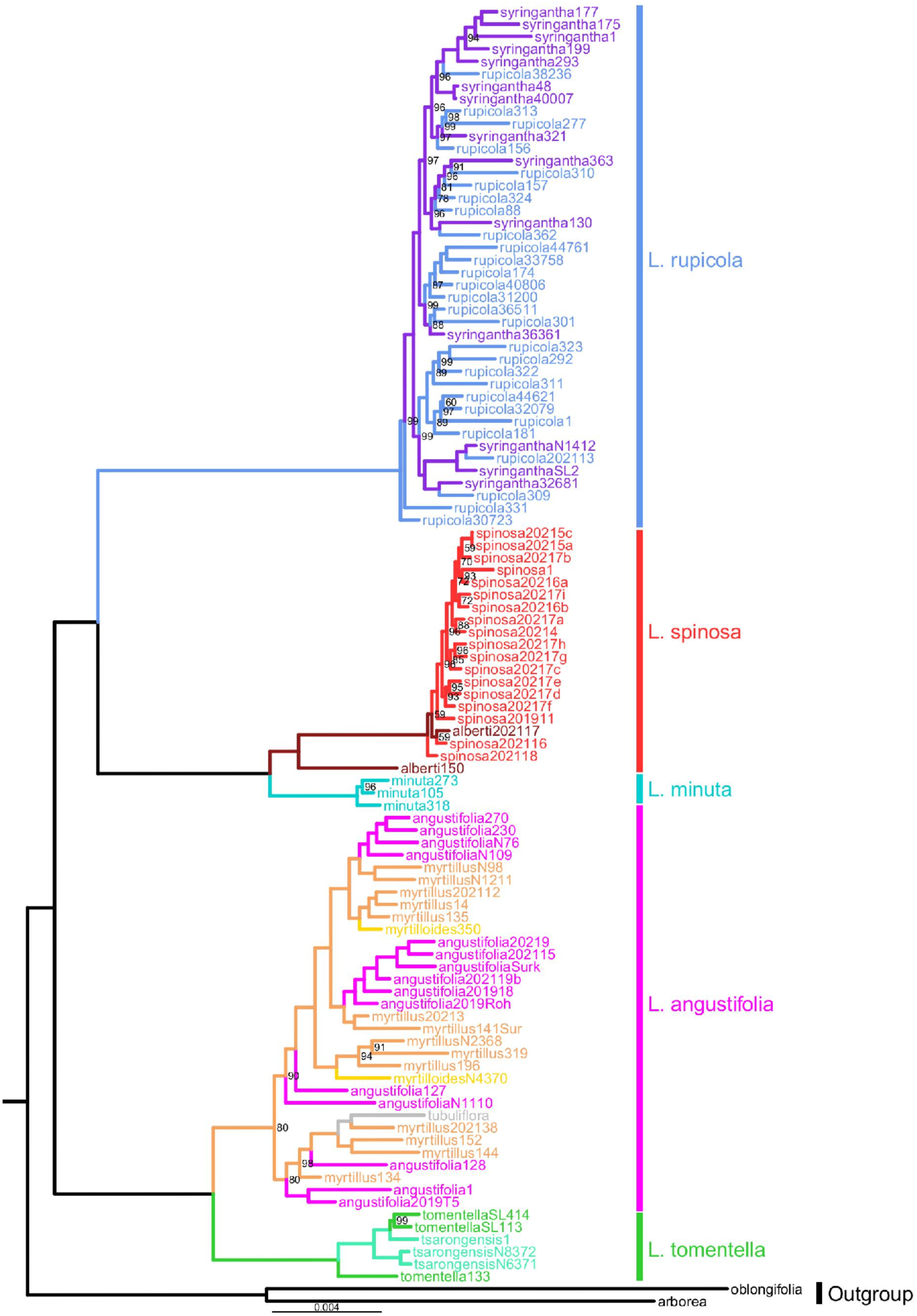
IQTree ML phylogram of *Isoxylosteum* estimated from RADseq loci assembled at a cluster threshold value of 90% and minimum number of samples per locus of 4. Bootstrap support values of 100% are not shown. Branch and tip label colors indicate previously

*L. rupicola* var. *rupicola* and *L. rupicola* var. *syringantha* samples form a clade, within which they are interdigitated rather than forming separate subclades. Similarly, within *L. angustifolia*, var. *angustifolia,* var. *myrtilloides,* and var. *myrtillus* do not form subclades, nor do the varieties of *L. tomentella* (var. *tomentella,* and var. *tsarongensis*). Likewise, *L. alberti* samples interdigitate with *L. spinosa* populations. The single sample of *L. tubuliflora* from Sikkim was found to be nested within the *L. angutifolia* clade. In contrast, the three sampled populations of *L. rupicola* var. *minuta* formed a distinct, well-supported clade.

IQ-Tree topologies were highly congruent across assemblies as were the Tetrad topologies, with only a few accessions changing positions within the *L. rupicola* and *L. spinosa* clades. Both methods recovered similar subclades that sort geographically, as noted below, with minor differences in accession placements within the species-level clades (Fig. S1).

Relationships within *L. tomentella* were identical across inference methods and assemblies. The Sikkim populations (SL414, SL113, and P1) form a clade sister to those from Nepal (N8372 and N6371), and together these are sister to the only sample from Bhutan (133; Fig. 2).

Within *L. angustifolia*, two major clades were recovered across inference methods — an eastern Himalayan clade, and a western Himalayan-central Himalayan clade (Fig. 2). Most populations of the western Himalayan-central Himalayan clade sort into two subclades: one western and one central Himalayan. Tree inference methods differed with respect to the placement of four central Himalayan populations (N4370, 196, N2368, and 319). While in IQTrees they formed a clade sister to the western Himalayan-central Himalayan clade, they were not recovered as a clade in Tetrad trees and were instead embedded in different parts of the western Himalayan-central Himalayan clade.

In *L. spinosa*, the only sample from Central Asia assigned at the outset to *L. alberti* (150) is sister to a western Himalayan clade (Fig. 2). Within the western Himalayan clade, the IQ-TREE and Tetrad phylogenies differ in the inferred relationships between the Spiti and Ladakh populations. IQ-Tree resolved an accession from Spiti (202118) as sister to a clade containing the other two Spiti populations (202116 and 202117) and another comprising all Ladakh populations. With Tetrad, Spiti and Ladakh populations form their own clades. Within the Ladakh clade, both methods recover two similar clades, a small clade comprising the southernmost populations and a larger clade containing the northern populations.

Within *L. rupicola*, all accessions were nested within one to two populations from southernmost Hengduan (30723 and 331; Fig. 2). Within the large clade, IQTree topologies recovered two major geographically distinct clades: (1) a clade spanning Himalaya, Sichuan, recognized species and varieties (see Fig. 1). The five species recognized here are labelled to the right. and southeastern Qinghai, and (2) a northern Sichuan clade extending into adjoining parts of Qinghai and Gansu. Clade (1) further sorted into a Himalayan-southern Sichuan clade and a Sichuan-southeastern Qinghai clade. Clade (2) contains two main subclades, one occupying northwestern Sichuan and adjoining parts of Qinghai and Tibet, and another spanning northern Sichuan and adjoining parts of southern Gansu and northeastern Qinghai. We note that while many of these clades were recovered in the Tetrad trees, they showed little resolution within *L. rupicola*. In these trees all populations are related to a sample from southernmost Hengduan (30723) and there is a Himalayan-southern Sichuan clade. However, none of the other details of the IQTree were recovered.

### Population Structure

STRUCTURE identified three clusters (K=3; Fig. 3A) which corresponded to: 1) *L. rupicola*, 2) *L. angustifolia-L. tomentella-L. tubuliflora*, and 3) *L. spinosa-L. alberti-L. minuta*, with *L. minuta* showing signs of admixture from *L. spinosa* and *L. rupicola*. When we ran further analyses on the *L. angustifolia-L. tomentella-L. tubuliflora* cluster, the preferred value of K=2 split the individuals into two distinct groups, corresponding to *L. angustifolia* and *L. tomentella*. The single sample of *L. tubuliflora* was recovered within the *L. angustifolia* cluster. Analyses focused on the *L. spinosa–L. alberti-L. minuta* cluster also favored K = 2, corresponding to *L. spinosa* and *L. minuta*. The two samples of *L. alberti* were recovered within the *L. spinosa* cluster. However, the Central Asian population of *L. alberti* (150) exhibited much greater genetic similarity to *L. minuta* than to the other *L. spinosa* samples. Both PCA approaches sorted the alleles into five clades corresponding to those inferred in the phylogeny (Figs. 3B, S2). The separation between *L. minuta* and *L. spinosa* was found to be smaller as compared to that between *L. minuta* and *L. rupicola*. Here again, the Central Asian accession of *L. alberti* (150) is closer to the *L. minuta* cluster than to other *L. spinosa* accessions.

**Figure 3.**
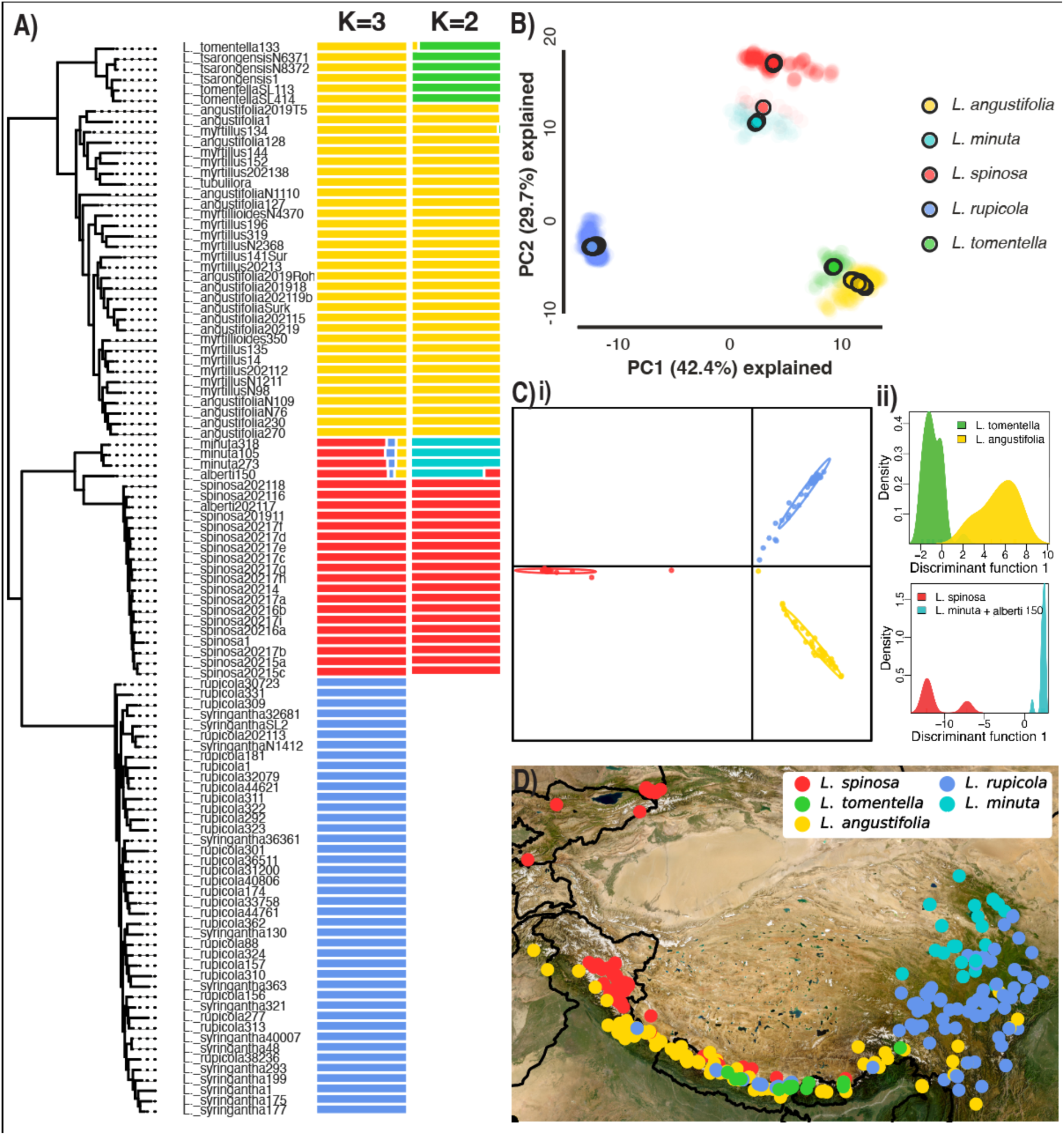
A) STRUCTURE analyses for *Isoxylosteum* and hierarchical clustering for *L. angustifolia-L. tomentella*, and *L. spinosa-L. minuta*. B) PCA of genetic variance of *Isoxylosteum* using K means of five. C) Results of DAPC analyses showing i) scatterplot for *Isoxylosteum*; the three recovered clusters correspond to *L. spinosa-L. minuta*, *L. tomentella-L. angustifolia*, and *L. rupicola*, and ii) density plots illustrating distribution of individuals along the first discriminant function for *L. spinosa-L. minuta*, and *L. tomentella-L. angustifolia* clades. Within the *L. spinosa-L. minuta* clade, the *L. alberti*150 sample was recovered within the *L. minuta* cluster. D) Map showing distribution of the five species recognized here.

DAPC favored the same three clusters within *Isoxylosteum*, and similar splits within each of the nested clusters, as recovered in STRUCTURE (Figs. 3Ci,ii). For the *Isoxylosteum*-wide dataset, a-score supported retention of 6 PCs. For the *L. angustifolia-L. tomentella-L. tubuliflora* and *L. spinosa–L. minuta* clusters, a-score supported retaining 1 PC each. Additionally, DAPC recovered *alberti150* within the *L. minuta* cluster (Fig 3Cii)

Weighted pairwise F_ST_ values between the five species reflect strongly structured, well-differentiated populations (Table S1). All pairwise comparisons yielded values greater than 0.49.

All combinations and algorithms of BPP analyses A10 and A11 supported the five species with high posterior probabilities (>98%). A11 analyses recovered the same species relationships as those inferred by the phylogeny, with high posterior probabilities (∼100%).

### Morphological Analyses

#### Leaves

Leaf length, leaf width, and leaf length-to-width ratio differed significantly among species (all P < 0.001; Table S2; Fig. S3). *L. tomentella* had the largest leaves (mean = 1.95 × 0.96 cm), followed by *L. rupicola* and *L. angustifolia*, whose leaves were approximately half as large on average. *L. spinosa* and *L. minuta* had the smallest leaves (1 × 0.15 cm, and 0.83 × 0.2 cm respectively; Fig. S3). *L. spinosa* had the highest leaf length-to-width ratio (∼7), followed by *L. minuta* (∼4) and *L. rupicola* (∼3.5). whereas *L. angustifolia* (∼3) and *L. tomentella* (∼2) had progressively lower ratios.

LDA based on leaf dimensions showed partial separation among species (Fig. S4). The first discriminant axis accounted for ∼98.5% of the among-group variation and was driven primarily by leaf width. Consistent with the LDA, leaf width exhibited greater interspecific differentiation than leaf length, differing significantly in 8 of the 10 species pairs compared with 6 pairs for leaf length (Table S3). Leave-one-out cross-validation correctly classified ∼59% of specimens (Table S4), exceeding the proportional chance criterion (26% based on observed species frequencies). Classification success was highest for *L. spinosa* (88%), followed by *L. angustifolia* (78%) and *L. tomentella* (59%). In contrast, fewer *L. rupicola* individuals were correctly classified (22%), while none of the *L. minuta* individuals were. *L. minuta* specimens were assigned to multiple species.

#### Flowers

PCA showed partial separation among the five recognized species (Fig. 4A). The sister species *L. angustifolia* and *L. tomentella* were differentiated primarily along PC1 which mostly reflects corolla traits (Figs. 4A, 4Ci; Table S5), whereas *L. spinosa* and *L. rupicola* were separated mainly along PC2 which mainly represents variation in ovary dimensions, and corolla and calyx tube widths (Table S5). The four samples of *L. minuta* mainly fell within the morphospace occupied by *L. rupicola* (Fig. 4A).

**Figure 4.**
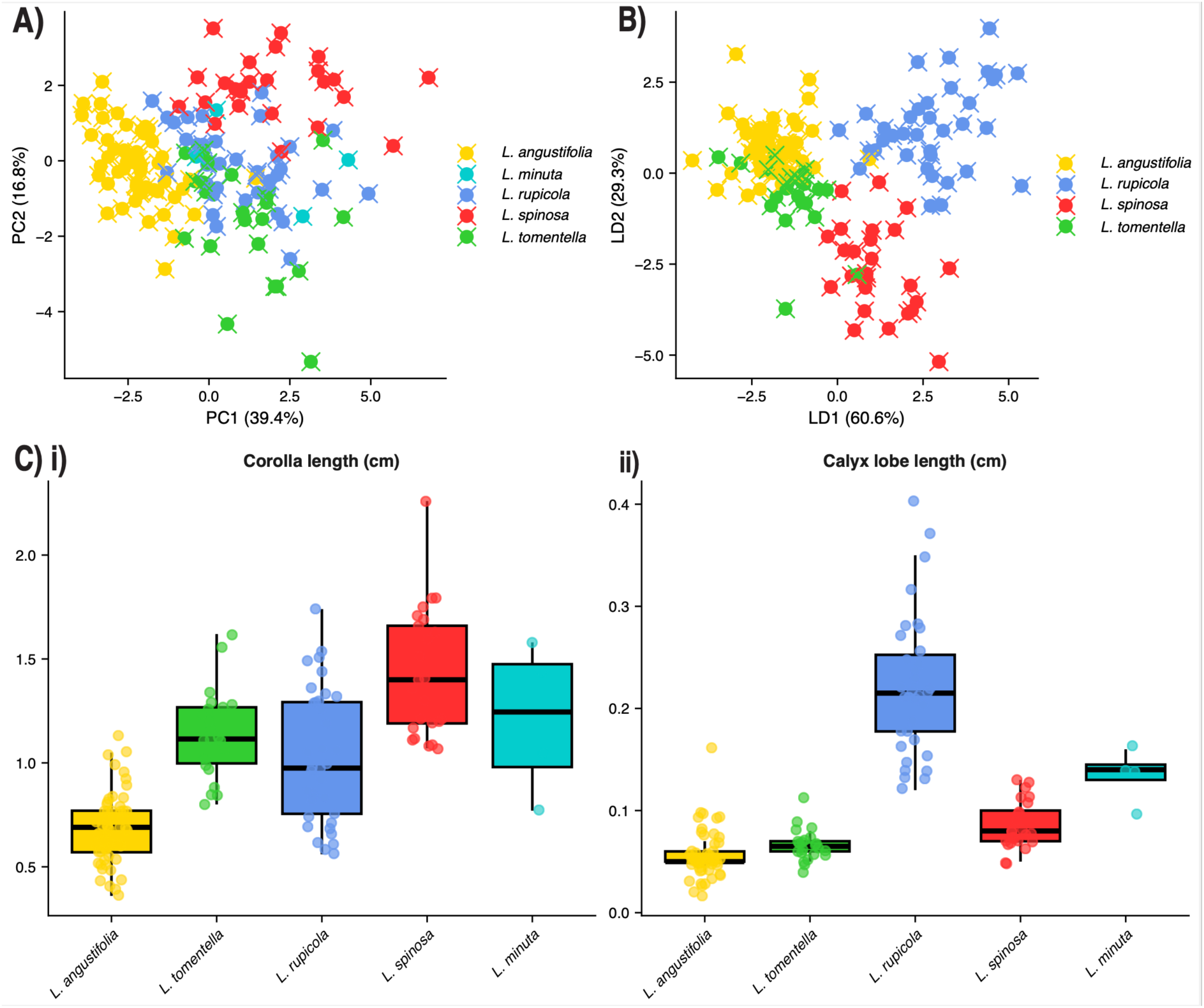
A) PCA of floral traits showing partial separation among species. B) LDA analyses performed on *L. spinosa, L. rupicola*, *L. tomentella*, and *L. angustifolia*. C) Boxplots showing variation among the five delimited species in i) corolla length, and ii) calyx lobe length, the latter of which had the highest discriminatory power among all traits. For boxplots of other floral traits, see Figure S5.

Despite partial overlaps in PC space, floral traits showed strong discriminatory power among *L. spinosa*, *L. rupicola*, *L. tomentella*, and *L. angustifolia* (Fig. 4B). LDA correctly assigned ∼89% of flowers to correct species in leave-one-out cross-validation. Classification accuracy was 88% or higher for all species except *L. tomentella* which was less reliably classified (∼68%), with most misclassifications occurring as *L. angustifolia* (Table S6A). The first two discriminant axes were primarily associated with PC3 and PC4, which captured variation in peduncle length, and calyx and ovary dimensions (Table S6B).

All 13 floral traits showed significant variation among species (p < 0.001; Figs. 4, S5; Table S7). Calyx lobe length showed the strongest discriminatory power, differing significantly in 8 of 10 species pairs (Fig. 4Cii; Table S8A). Among species, *L. angustifolia* exhibited the greatest number of significant pairwise differences, differing from *L. rupicola* in 12 traits and from *L. spinosa* and *L. tomentella* in 10 traits each (Table S8B).

### Ancestral state reconstructions

Our findings on morphological character evolution in *Isoxylosteum* are summarized in Figure 5, which is based on maximum likelihood reconstructions of 11 traits inferred across all of *Lonicera* (Figure S6). The ancestor of *Isoxylosteum* is inferred to have lacked thorns and borne two glabrous leaves per node with prominent secondary veins (Fig. 5). Radially symmetrical flowers evolved in the ancestor of *Isoxylosteum* (Figs. 5, 6A-E, S6i), likely bearing five lobes, a glabrous corolla tube, and stigma and stamens exserted. We note, however, that the latter reconstruction is only marginally supported over the deeply inserted stigma and stamen condition (Fig. S6ix). Finally, the common ancestor also likely had bracteoles fused into a partial cupule (Fig. 6C) and free ovaries (Figs. 6M-O) that matured into red fruits.

**Figure 5.**
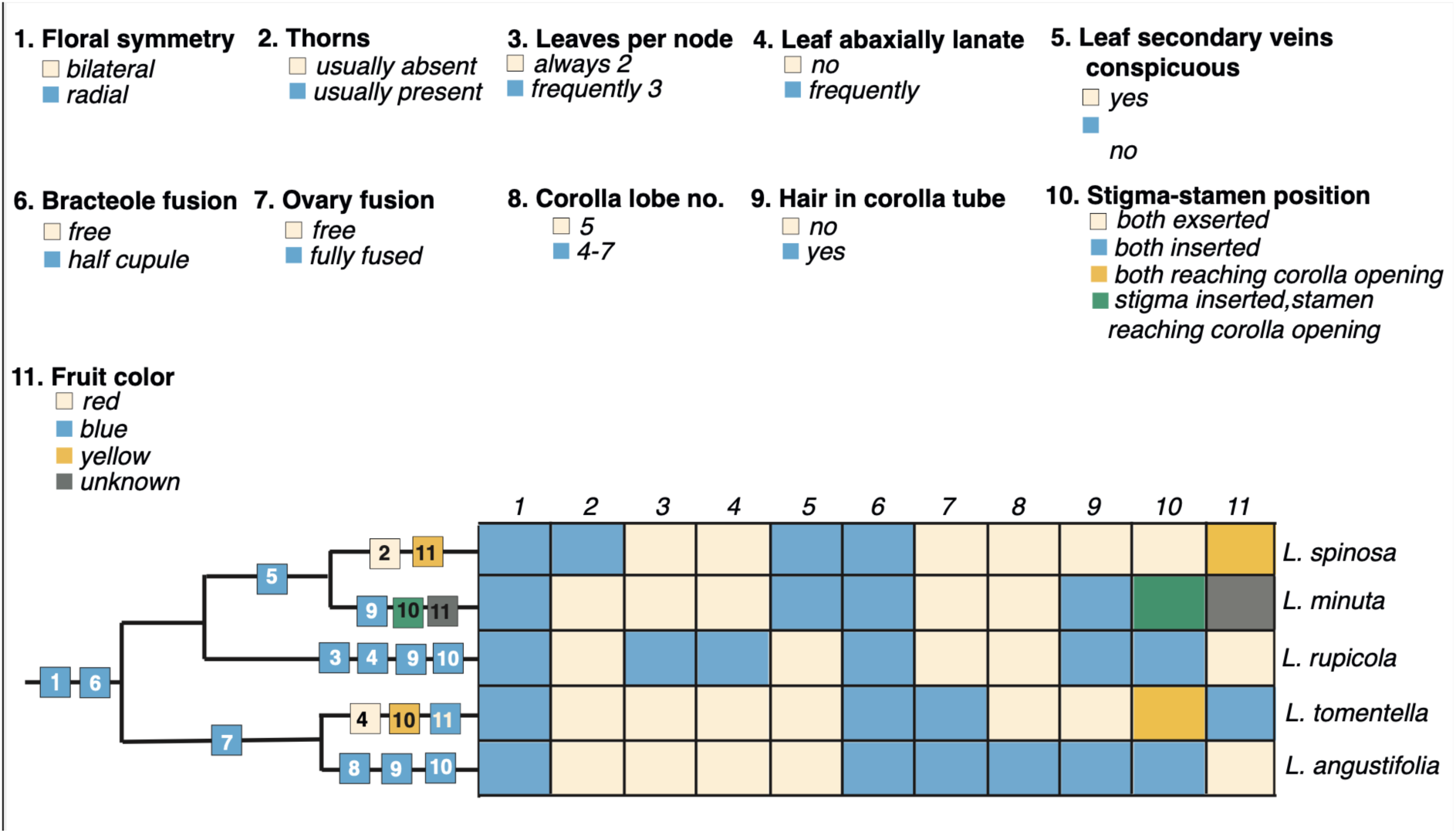
A summary of morphological character evolution in *Isoxylosteum* (note that *L. tomentella* produces tomentose leaves frequently). Reconstructions for each individual character across *Lonicera* are presented in Figure S6.

**Figure 6.**
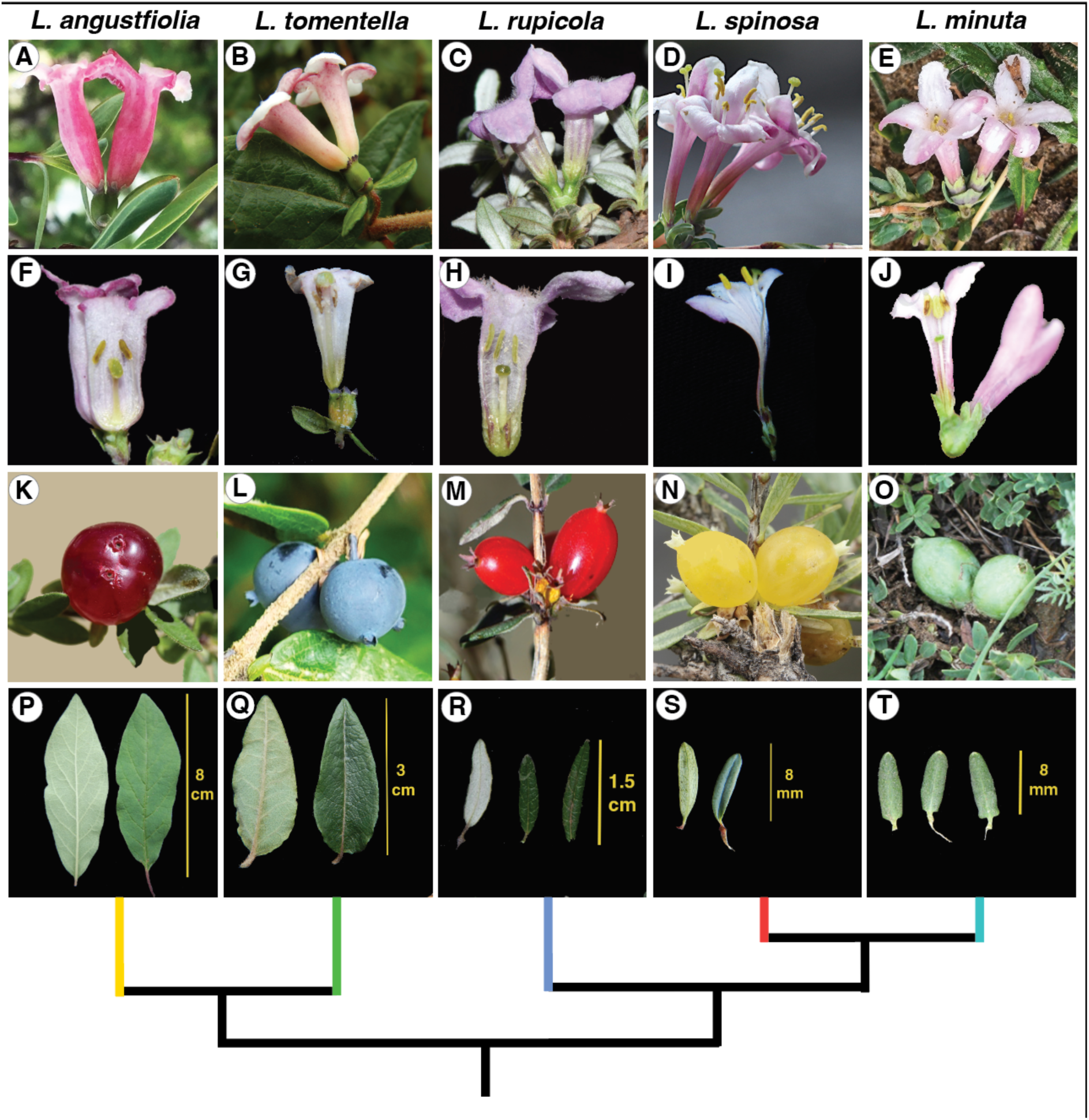
Diversity of morphological traits across the five delimited species of *Isoxylosteum*. Top row: flowers showing radial symmetry, partially fused bracteoles, and variation in floral traits; second row: longitudinal sections of flowers showing the diversity in stamen and stigma length across the clade; third row: mature ovaries illustrating fruit color diversity and clade-wise differences in ovary fusion (free versus completely fused); fourth row: leaves showing variation in size, shape, secondary vein visibility, and pubescence. (Photos courtesy: A, © Wangtianming, some rights reserved (CC-BY), via iNaturalist Observation 369687827; C, © Zhu Xinxin, rights reserved, via Plant Photo Bank of China (https://ppbc.iplant.cn/)

There are clear-cut synapomorphies uniting each of the major clades within *Isoxylosteum* (Fig. 5). *L. tomentella* and *L. angustifolia* share fully fused ovaries (Figs. 6K, L), while the *L. rupicola-L. minuta-L. spinosa* clade is characterized by free ovaries (Figs. 6M-O). *L. minuta* and *L. spinosa* share thin leaves with inconspicuous secondary veins (Figs. 6S, T). Each species also exhibits distinct autapomorphies. For example, *L. rupicola* often bears three leaves per node, *L. angustifolia* frequently has four to seven corolla lobes, *L. spinosa* produces thorns and yellow fruits (Fig. 6N), and *L. tomentella* bears blue fruits (Fig. 6L). Corolla pubescence and stigma-anther position show homoplasy (Figs. 6F-J). Pubescent corolla tubes are inferred to have evolved three times independently, in *L. minuta, L. rupicola,* and *L. angustifolia*, and the combination of inserted stamens and stigmas likely evolved independently in *L. rupicola* and *L. angustifolia* (Figs. 5, 6, S6ix). A dichotomous key to the fives species is provided in Appendix S2.

### Distribution and altitudinal ranges

*Isoxylosteum* is endemic to the Himalayan, Hengduan, Pamiro-Alai and Tien Shan regions and, with populations of *L. spinosa* to the north, it forms an incomplete ring around the Tibetan plateau (Fig. 3D). *L. spinosa* is predominantly distributed in the western Himalaya, but extends around the north of Tibet to Central Asia and around the south of Tibet to central Himalaya. It occupies the highest elevational ranges in *Isoxylosteum*, from ∼3200 to ∼4800 m. *L. rupicola’s* distribution is centred in the Hengduan region, occurring from ∼2,050 to ∼5,150 m; it tapers off westward toward the junction of central and western Himalaya. *L. tomentella* is narrowly distributed in central and eastern Himalaya from ∼2000 to ∼4540 m. *L. angustifolia* spans the entire Himalayan range from west to east, from ∼2000 to ∼4400 m, extending into the adjacent parts of Hengduan. The range of *L. minuta* is the narrowest—it is restricted almost completely to eastern Qinghai and adjoining parts of Gansu and Sichuan from ∼3000-∼4400 m.

## Discussion

### Species delimitation

We view species as independently evolving metapopulation lineages (de Queiroz 2007, 1998; Donoghue 2022), where such independence is judged by evaluating various lines of evidence that have in the past been equated with species criteria under different species concepts (e.g., reproductive isolation under the biological species concept; Mayr, 1942). Here we interpret all of the evidence we have presented as supporting the recognition of five species: *L. tomentella*, *L. angustifolia, L. rupicola, L. spinosa*, and *L. minuta.* These are reciprocally monophyletic based on our RADseq data and they show distinct patterns of genetic diversity in our STRUCTURE, DAPC, PCA, and BPP analyses. Each of these lineages also has a specifiable geographic range and ecological niche (judged here by habitat type and elevational range). These species are also distinguished by strongly differentiated floral traits and moderately differentiated leaf traits. Finally, they are each marked by evolutionarily derived floral traits that suggest that shifts to different pollinators may have promoted their reproductive isolation.

The previously recognized varieties within *L. angustifolia* (var. *angustifolia*, var. *myrtillus*, and var. *myrtilloides*), two of the *L. rupicola* varieties (var. *rupicola*, and var. *syringantha*), and the two *L. tomentella* varieties (var. *tomentella*, and var. *tsarongensis*) are not being recognized here, and neither is *L. alberti.* Our combined evidence does not support their status as independently evolving lineages. They do not form clades in our RADseq trees, nor distinct clusters in our PCA, STRUCTURE, DAPC, or BPP analyses. In some cases, they differ morphologically, but several traits appear to have arisen independently in relation to their occupation of different habitats at different elevations (see below). *L. rupicola* var. *minuta*, on the other hand, forms a distinct lineage that is most closely related to *L. spinosa*, and it is here recognized as a separate species. The closer relationship of *L. minuta* to *L. spinosa* than to *L. rupicola* is strongly supported by our phylogenetic and population analyses as well as morphological synapomorphies and ecological similarities as discussed below. The separation of *L. minuta* from *L. spinosa* is supported by BPP analyses and a high pairwise F_ST_ value of 0.66, which is even higher than that of the traditionally recognized species pair, *L. angustifolia* and *L. tomentella* (0.49; Table S1). The status of *L. tubuliflora* remains uncertain as discussed below.

### Morphological Variation and Evolution

Having recognized the existence of five species of *Isoxylosteum*, we now consider morphological diversity in the clade in greater detail, with an emphasis on variation that has figured in past taxonomic decisions (Fig. 6). We especially highlight parallel morphological shifts within species as sources of confusion, but also call attention to potential targets for more detailed functional studies.

#### Plant habit

*soxylosteum* species are all shrubs, but they differ dramatically in size, even within a species. *L. tomentella* grows up to 4-5 m but may be less than 1 m tall at higher elevations. Its sister species, *L. angustifolia*, can grow nearly as tall but at high elevations it can be as short as 3 cm. In the field we encountered this dramatic variation within a span of ∼1 km. The dwarf and intermediate forms (var. *myrtillus* and var. *myrtilloides*, respectively) are nested within lineages of taller plants (var. *angustifolia*) in the major subclades (Fig. 2). The three varieties were found to sort geographically and elevationally: var. *angustifolia* occupies the outer Himalayan ranges and adjacent Hengduan regions at lower elevations (∼2000–∼3950 m), var. *myrtilloides* occurs primarily in the inner Himalayan and the adjacent Hengduan ranges at intermediate elevations (∼2760–3950 m), while var. *myrtillus* occupies the northernmost and innermost Himalayan regions at the highest elevations (∼2900–4400 m). This suggests parallel local adaptation to alpine environments in *L. angustifolia* (Wos et al. 2022). It is unclear whether this has a genetic underpinning or is simply a plastic response. *L. rupicola* can grow to 2.5 m but it too can express an extreme dwarf form of only ∼3 cm. *L. spinosa* and *L. minuta* plants are the most consistently short in stature, often reaching ∼30 cm, but we have also seen plants of *L. spinosa* up to ∼0.7 m in height.

Despite its name, *L. spinosa* does not bear spines but thorns. *L. rupicola* and *L. angustifolia* can also produce thorns in high-elevation, dry environments, making it hard to differentiate between the herbarium specimens of *L. spinosa* and the reduced forms of *L. rupicola* and *L. angustifolia* in the absence of reproductive traits. *L. alberti* was separated from *L. spinosa* mainly on the basis that it does not bear thorns but we found them to be genetically interdigitated.

#### Leaf traits

All *Isoxylosteum* species have two opposite leaves but *L. rupicola* frequently bears leaves in whorls of three (Fig. 5). Together with *L. tomentella*, it also differs from other species in frequently bearing pubescence. Its leaves are often covered with white pubescence on the lower leaf surfaces. (Fig. 6R). Although the recognition of *L. rupicola* varieties has been based on this pubescence trait (Rehder 1903), in the field we observed considerable variation within individual plants. Rehder (1903) also noted that, in the form var. *rupicola*, the flowering branches can have abaxially villous leaves while the rest are glabrous. Interestingly, these varieties show geographic structuring within the Hengduan mountains: the glabrous variety *syringantha* occurs predominantly in the eastern part of the range while the pubescent variety *rupicola* occupies more western areas (Fig. 1). This could reflect adaptation to contrasting environmental conditions across the Hengduan region, such as differences in temperature, precipitation, and other climatic variables (Johnson 1975; Bickford, 2016). Whether these morphological differences are associated with environmental divergence remains to be tested using ecological and environmental niche analyses.

*L. tomentella* leaves and stems are covered by a soft brown-orange pubescence. Its two varieties were differentiated on the basis of leaf pubescence, number of lateral veins, and whether the lower surface is glaucous. Those with less hairy, glaucous leaves bearing 4-5 pairs of lateral veins were assigned to var. *tsarongensis*, while those with more pubescent, non-glaucous leaves and 7-8 pairs of lateral veins were recognized as var. *tomentella*. We found the two varieties to be genetically intermixed. Moreover, most specimens could not be neatly binned into the two varieties—even though many specimens with glaucous leaves were less pubescent, we found glaucous-looking specimens with considerable pubescence, bearing lateral veins ranging anywhere from four to more than eight pairs.

Leaf size and proportions broadly corresponded with elevational distribution, with the smallest and narrowest leaves occurring in the highest-elevation species, *L. spinosa* and *L. minuta*, and the largest and broadest leaves in *L. tomentella*, which occupies relatively lower elevations (Fig. S3). This pattern is consistent with the generally smaller leaf sizes observed in alpine environments, where reduced leaf area may help limit water loss and mitigate exposure to cold, dry, and windy conditions (Körner, 2003).

*Isoxylosteum* species also vary in leaf blade thickness: *L. rupicola* and *L. tomentella* have thick, leathery leaves while those of the others are thinner (Fig. 6P-T). Additionally, all species have entire leaf margins but *L. angustifolia* leaf margins can be sinuate. In *L. minuta* and *L. spinosa*, the secondary veins are inconspicuous whereas those in other species are clearly visible.

*Isoxylosteum* species also exhibit variation in leaf blade, apex, and base shape. Those occupying the highest elevations have narrowly shaped leaves: *L. spinosa* has linear leaves with an acute apex and an attenuate base, and *L. minuta* has lanceolate-linear to acute leaves with acute-obtuse apex and cuneate-attenuate base. The lower elevation *L. tomentella* bears elliptic to ovate leaves, with base cuneate, rounded, truncate or cordate, and apex acute to obtuse. Species occupying the widest elevational ranges show the most varied leaf shapes. *L. rupicola* leaves can be linear, lanceolate, elliptic, or ovate, with base cuneate, rounded, truncate to cordate, and apex acute to acuminate. Similarly, *L. angustifolia* leaves can be elliptic, ovate, rhomboidal, obovate, linear, lanceolate, with base ranging from cuneate to attenuate, and apex generally acute. Notably, *L. angustifolia* leaf, apex and base shape can observation 3671614; D, © Tibetissimo, some rights reserved (CC-BY-NC), D via iNaturalist Observation 374960480; E, © niuzhi_li, some rights reserved (CC-BY-NC), via iNaturalist Observation 371965298; F, © Zhu Xinxin, rights reserved, via Plant Photo Bank of China Observation 3671516; H, © Zhu Xinxin, rights reserved, via Plant Photo Bank of China Observation 3671620; J, © Zhu Xinxin, rights reserved, via Plant Photo Bank of China Observation 6204743; K, © Shen Zhuomin, rights reserved, via Plant Photo Bank of China Observation 23623534; M, © Zhu Renbin, rights reserved, via Plant Photo Bank of China Observation 5923806; N, © Zhao Xinjie, rights reserved, via Plant Photo Bank of China Observation 24643889; O, © Liu Xiang, rights reserved, via Plant Photo Bank of China Observation 4778089; S, © Zhu Xinxin, rights reserved, via Plant Photo Bank of China Observation 3671680; T, © Zhu Xinxin, rights reserved, via Photo Bank of China Observation 6204754).

vary widely even among individuals growing in close proximity and even among leaves on the same branch. Given the highly variable habit and leaf morphology of *L. angustifolia*, it is not surprising that we found its herbarium specimens frequently misidentified as one of the other *Isoxylosteum* species, especially in the absence of reproductive traits. Yet, surprisingly it had a high classification success based on leaf dimensions alone (78%; Table S4).

#### Floral Traits

*soxylosteum* species are united by having pink to white radially symmetrical corollas (Fig. 6A-E) with five nectaries. All species have five corolla lobes, but *L. angustifolia* has frequently been reported to bear four to seven lobes (Rehder 1903; Yang et al. 2011). In the field we observed adjacent flowers of *L. angustifolia* fusing to varying degrees. Many populations had corollas fused along the sides of the tube facing one another, while the flowers otherwise maintained their individual identity. However, in some populations we observed the complete fusion of adjacent flowers, with the resulting mega-flower bearing a two-headed stigma, several stamens, and seven to eight corolla lobes. This fusion phenomenon appears to account for reports of more than five corolla lobes in *L. angustifolia*.

All species possess bracteoles united into a partial cupule around the two ovaries (Fig. 6A-E). This structure is often lobed and can vary from a small cup at the base, as in *L. tomentella*, to nearly covering the ovaries at young stages in some populations of *L. angustifolia*.

In keeping with the compact architecture of cold-adapted plants, species occupying the highest elevations (*L. spinosa*-*L. minuta*-*L. rupicola* clade) bear shorter peduncles (Fig. S5). *L. rupicola-L. spinosa-L. minuta* clade generally bears longer corollas, wider corolla diameters, and bigger corolla and calyx lobes but slightly narrower corolla tubes than *L. angustifolia-L. tomentella* clade. Calyx tube width at the base and ovary length do not sort clade-wise, but intriguingly, they show a similar pattern*: L. tomentella* has the highest values followed by *L. rupicola*, whereas *L. angustifolia* has the smallest values (Fig. S5).

Plants in the *L. tomentella-L. angustifolia* clade have fully fused ovaries (Figs. 6 K,L), while the ancestral condition of having free ovaries has been retained in the other clade (Figs. 6M-O). In keeping with that, *L. angustifolia-L. tomentella* clade was found to bear wider ovaries but *L. angustifolia* ovaries were found to be only slightly wider than the species with free ovaries (mean 1.4 cm versus 1.2 cm; Fig. S5). As in multiple traits, *L. angustifolia* can be variable in ovary fusion, and we have observed only partially fused and even free ovaries in some cases which could account for this. This species would be an excellent candidate for detailed studies of organ fusion in *Lonicera* (see Srivastav et al. 2023).

Interestingly, although corolla traits together with ovary dimensions and calyx tube widths accounted for much of the overall floral variation, species discrimination depended more strongly on variation in peduncle, calyx, and ovary traits (Tables S5, S6B). Despite contributing relatively little to the major axes of overall variation, calyx lobe length emerged as the most informative trait for species delimitation (Table S8A). One possible explanation is that corolla dimensions could be more evolutionarily labile because they are shaped by pollinator-mediated selection (Wu et al., 2018) whereas calyx and ovary traits may be under weaker or different selective pressures in this case.

The variable relative position of the stamens and the stigma is an underappreciated trait within *Isoxylosteum* (Fig. 6F-J; Fig. 7). In addition to having short corollas, *L. angustifolia* and *L. rupicola* appear to have independently evolved short, stout styles, with the stigma and stamens positioned well within the corolla tube. Consequently, together with the dense hairs in the corolla throat, these reproductive structures are not visible from outside the flower. This stamen-stigma length is a unique combination across all of *Lonicera* (Fig. S6ix). In *L. minuta* the stigma is situated within the tube, but the stamens reach the level of the opening of the corolla (Fig. 6J). It also bears hair in the corolla throat but these are less dense than those in *L. angustifolia* and *L. rupicola*. In *L. tomentella* both the stamens and the stigma reach the corolla opening (Fig. 6G). In contrast to the others, *L. spinosa* has a long, slender style bearing completely exserted stigma, as well as exserted stamens with thin filaments (Fig. 6I). The *L. spinosa* condition is found across all other ∼135 *Lonicera* species outside *Isoxylosteum*, except one species (*L. mexicana*). This exserted stigma-stamen position is marginally favored as the ancestral condition for *Isoxylosteum* over completely inserted stigma-stamen (Figs. 7, S6ix).

**Figure 7.**
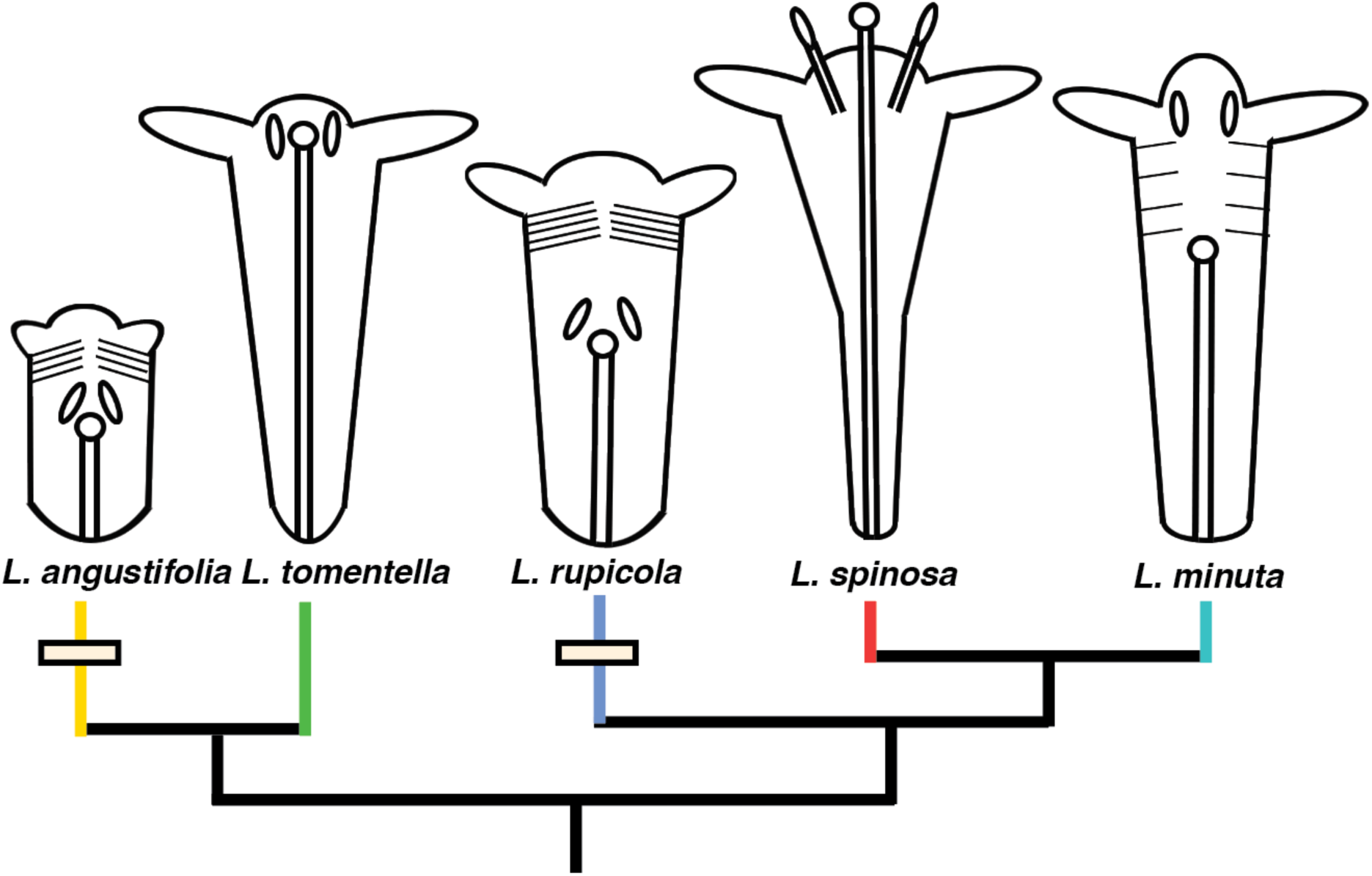
*Isoxylosteum* phylogeny with schematic drawings of longitudinal sections of flowers that illustrate the position of the stigma and stamens in relation to the corolla tube. (only two of the five stamens shown; flower parts scaled to reflect actual size differences). The small lines within the corolla tube depict pubescence in *L. angustifolia* and *L. rupicola*, and less densely in *L. minuta*. Cream bars mark the independent evolution of deeply embedded stigmas and anthers in *L. rupicola* and *L. angustifolia*. When measured at the widest point, its corolla tube is second widest, after *L. tomentella* (Fig. S5). This widening towards the corolla mouth may facilitate accommodation of the exceptionally long stamens and stigma of *L. spinosa* within the floral bud.

*L. angustifolia* bears the smallest flowers overall, although its corolla tube and ovary are among the widest (Fig. S5). *L. minuta* is characterized by the longest calyx tube and the widest calyx lobes. *L. spinosa* has the longest and widest corolla diameter. However, its corolla tube is the narrowest at the midpoint. Interestingly, whereas the corolla tubes of the other species remain relatively uniform in width along their length or widen only gradually toward the opening, that of *L. spinosa* widens markedly from the midpoint toward the mouth.

Detailed comparative studies of pollination biology are clearly called for, but our presumption is that changes in flower architecture have been accompanied by shifts in the predominant pollinating insects. The likely derived flower form in *L. angustifolia* and *L. rupicola* may relate to pollination by long-tongued flies and the bumblebees that are so diverse in this region (Williams 1998, Williams et al. 2009).

#### Fruit Traits

Mature fruit color varies dramatically in *Isoxylosteum*. *L. angustifolia* has red fruits, which we infer to be the ancestral condition (Figs. 6K, S6x), and, similarly, *L. rupicola* has red to reddish-orange fruits (Fig. 6M). In contrast, *L. tomentella* has derived blue-black fruits with a glaucous bloom (Fig. 6L), and *L. spinosa* has translucent yellow fruits that turn yellowish-orange and dry out as they mature ((Fig. 6M). In the field, we observed that these fruits turn orange and developed a toffee-like consistency at maturity, after which they fell on or near the parent plant and withered away, gradually exposing the seeds. We note that *L. spinosa* fruit color has been erroneously described as white or pale violet in most taxonomic treatments (Rehder 1903;Yang et al. 2011). The fruit color of *L. minuta* remains uncertain.

### Status of L. tubuliflora and L. minuta

The species status of *L. tubuliflora* remains uncertain. Based on our analyses it would appear to be nested well within *L. angustifolia* (Figs. 2, 3). This is at odds with the view of Rehder (1913) that it is most closely aligned with *L. rupicola*, but it makes better sense in terms of morphology, as also noted by Yang et al. (2011). *L. tubuliflora* was more similar to *L. angustifolia* in quantitative floral traits, differing significantly in only two measured traits (corolla length and corolla tube length), compared with four traits for *L. rupicola* (Table S9). In floral PC space, the six samples of *L. tubuliflora* largely overlapped with *L. angustifolia* (Fig. S7)*. L. tubuliflora* was distinguished from *L. angustifolia* by its free ovaries, a narrower, tubular corolla tube with mouth slightly constricted near the opening, and shorter corolla lobes ∼1.5 mm (Yang et al. 2011), compared with partially to fully fused ovaries, tubular-campanulate corolla, and lobes generally longer than 1.5 mm. However, these diagnostic characters showed substantial overlap between the two taxa. The corolla width of *L. tubuliflora* was not found to be significantly narrower than that of *L. angustifolia* (mean 2.3 mm versus 2.8 mm). The mean corolla lobe length of *L. tubuliflora* was 1.5 mm, whereas *L. angustifolia* had even shorter lobes (mean 1.2 mm). Additionally, we observed *L. angustifolia* populations with free ovaries, both in the field and herbarium specimens, and its corolla shape varied from tubular-campanulate to narrowly tubular. Both species also produce red berries and have been observed to undergo a similar floral color change from white to yellow. Furthermore, our leaf LDA classified all 21 *L. tubuliflora* leaves as *L. angustifolia* (Table S10). However, corolla and corolla tube lengths of *L. tubuliflora* were found to be significantly longer (means 9.5 mm versus 6.9 mm; means 9.5 mm versus 6.8 mm), suggesting that it might be an overlooked difference. Conclusions regarding the status of *L. tubuliflora* remain preliminary because our molecular dataset included only a single specimen of this species, collected from Sikkim in the eastern Himalaya, outside its previously reported range in Sichuan, China. Resolving the status of *L. tubuliflora* will require the analyses of additional collections, particularly those collected within its presently known range.

*L. minuta* was considered to be a variety of *L. rupicola* but genetic analyses strongly support its closer relationship to *L. spinosa* (Figs. 2, 3). Both *L. minuta* and *L. spinosa* inhabit cold deserts while *L. rupicola* occurs in cold deserts, alpine environments as well as at lower elevations at the forest margins. Several morphological characters also lend support to the closer relationship of *L. minuta* to *L. spinosa*: they both have the narrowest leaves within *Isoxylosteum* (Fig. S3) and also share thin leaf blades with inconspicuous secondary veins, in contrast to the thick leaves and prominent secondary veins of *L. rupicola* (Fig. SR-T). *L. minuta* and *L. spinosa* also do not differ significantly in either leaf length or width, whereas *L. minuta* differs significantly from *L. rupicola* in both traits (Table S11). Among the floral characters, *L. minuta* differs significantly from *L. spinosa* in peduncle length, whereas it differs from *L. rupicola* in calyx lobe length (Table S8B). The strongest morphological distinction of *L. minuta* from these two species as well as from the other *Isoxylosteum* species, however, is the unique combination of stamen and style lengths, which, together with the genomic evidence, supports its recognition as a distinct species (Figs. 5, 6F-J). Fruit color will further clarify the status of *L. minuta* but we were not able to find any photos of its matured fruits. *L. minuta* grows in shifting sand dunes. It bears highly branched, largely subterranean stems that produce extremely short shoots annually (3-5 cm) since whichever part of the plant is not covered by the loess dies away in winters (Batalin 1892; Rehder 1903; Yang et al. 2011). The slow growth rate of this species along its very narrow distribution warrants conservation priority for *L. minuta*.

Finally, it is important to consider our accession *alberti150* and its bearing on species delimitation. As noted above, this sample from the northern rim of the Tibetan Plateau is genetically nearly identical to our *L. minuta* samples from the Qinghai region (Figs 1, 3), yet it is sister to *L. spinosa* specimens in our phylogenetic analyses. Two possibilities are consistent with this finding: (1) the existence of a series of intermediate populations spanning that region, or (2) admixture between *L. spinosa* and *L. minuta* in the past. Although admixture may seem unlikely given that *L. minuta* is not currently known from this region, we cannot exclude the possibility that its distribution has contracted since past contact, or that unsampled populations persist there. The only way to distinguish among these possibilities is through additional fieldwork in that region to find and sample such populations if they exist. In the meantime, pending further collecting, we think it best to exclude accession *alberti150* from either *L. spinosa* or *L. minuta*.

### A taxonomic foundation for future research

The recognition of five species within *Isoxylosteum* is strongly supported by the consilience of multiple lines of evidence. Each species forms a well-supported clade in our several phylogenetic and population analyses, with high genetic distances between them. They also differ in morphological traits and with respect to their geographic ranges and the habitats that they occupy. The agreement of all of these lines of evidence makes species delimitation in this group refreshingly straightforward. At the same time, these data provide compelling evidence against the recognition of the previously named varieties, with the exception of *L. rupicola* var. *minuta*, which we now recognize as a separate species. Our most surprising result is the finding that the former *L. rupicola* var. *minuta* is more closely related to the geographically widely separated *L. spinosa* than it is to any populations within *L. rupicola*.

The phylogenetic and species delimitation analyses presented here set the stage for biogeographic and niche evolution studies in *Isoxylosteum* and its bearing on the broader issue of the historical assembly of the high-altitude flora of this extraordinary mountain-plateau system (Srivastav et al., in prep.). Our analyses of the remarkable morphological variation found within and among *Isoxylosteum* species demonstrates that in many respects this small clade is a microcosm of trait diversity across *Lonicera* as a whole. As such, we have established a firm foundation for more detailed analyses of traits that have attracted attention more broadly, including the fusion of bracteoles and fruits (Srivastav et al., 2023). We have also identified likely cases of homoplasy in plant architecture and leaf traits in relation to shifts in elevation and habitat type, and in floral syndromes, likely in relation to shifts in pollination biology.

## Acknowledgements

This work was funded by National Science Foundation grants to M.J.D. (DEB-1929533), Wendy Clement (DEB-1929670), and Dianella Howarth (DEB-1929674), the Macmillan Centre at Yale University, and the Yale Institute for Biospheric Studies. We especially thank all the volunteers and field assistants who helped with collecting in different parts of the Himalaya, especially Neha Tiwari, Yudyang Kaushik, Dungzo, Lepe, Buddha, Birhang, and Krishna. We want to thank Biodiversity of the Hengduan Mountains project for providing samples for the study. We also acknowledge the Director and Dean of the Wildlife Institute of India for institutional support. Thanks also to members of the Donoghue, Near, Munoz, and Edwards labs at Yale for helpful discussions. Finally, we thank the Forest department of Nepal and the National Biodiversity Authority, India, and Principal Chief Conservator of Forests and Chief Wildlife Wardens of the Himalayan States for granting us collecting permits. We also want to thank the Photo Bank of China (PPBC) for allowing us to use their photos.

## Author Contributions

M.S. and M.J.D. conceptualized and designed the study. M.S. conducted field collections, lab work, analyses, figure preparation, and drafted the manuscript. A.K., and G.S.R. assisted with field collections, and acquiring collection permits. A.K., G.S.R, R.H.R., and M.J.D. advised throughout. All authors contributed to writing and editing the manuscript.

## Data availability

All sequences generated for this work are available on NCBI Sequence Read Archive (SRA) under the BioProject ##. Alignments, trees, and morphological character matrix are available on the Dryad Digital Repository (https://doi.org.#).

## APPENDIX 1

Collection and locality details of sampled specimens. The sources for herbaria and silica gel material are abbreviated as follows: Hengduan - Biodiversity of the Hengduan mountains project (http://hengduan.huh.harvard.edu/fieldnotes); WII – Wildlife Institute of India; KATH - National Herbarium and Plant Laboratories, Kathmandu; MW - Moscow State University Herbarium (https://plant.depo.msu.ru/); MO - The Missouri Botanical Garden’s Herbarium; YU - The Yale Herbarium; K - The Herbarium at the Royal Botanic Gardens Kew; E - Royal Botanic Garden Edinburgh Herbarium; F - The John G. Searle Herbarium, Chicago; KUN – herbarium at the Kunming Institute of Botany, Chinese Academy of Sciences herbarium; and MA - Real Jardín Botánico de Madrid herbarium.

| SN | Accession No. | Latitude | Longitude | Elevation<br>(in m) | Herbarium /Source | Herbarium<br>Voucher ID<br>/Collection No. |
| --- | --- | --- | --- | --- | --- | --- |
| 1 | Lonicera_rupicola1 | 33.124722 | 97.458889 | 4295 | Hengduan | 44775 |
| 2 | Lonicera_syringantha1 | 33.668611 | 103.242778 | 2880 | Hengduan | 40196 |
| 3 | Lonicera_rupicola36361 | 31.646944 | 100.266389 | 3545 | Hengduan | 36361 |
| 4 | Lonicera_rupicola44621 | 31.893611 | 96.596111 | 4100-4200 | Hengduan | 44621 |
| 5 | Lonicera_syringantha32681 | 28.96 | 102.103611 | 3800-4000 | Hengduan | 32681 |
| 6 | Lonicera_syringantha40007 | 33.108611 | 102.631944 | 3485-3520 | Hengduan | 40007 |
| 7 | Lonicera_rupicola31200 | 31.448333 | 98.165278 | 3625 | Hengduan | 31200 |
| 8 | Lonicera_rupicola38236 | 30.952778 | 102.296667 | 2903 | Hengduan | 38236 |
| 9 | Lonicera_rupicola44761 | 32.836111 | 95.5625 | 3980-4125 | Hengduan | 44761 |
| 10 | Lonicera_rupicola40806 | 30.849722 | 95.439167 | 3755 | Hengduan | 40806 |
| 11 | Lonicera_rupicola33758 | 32.479722 | 98.021944 | 3380 | Hengduan | 33758 |
| 12 | Lonicera_rupicola30723 | 29.1 | 99.692222 | 4000-4500 | Hengduan | 30723 |
| 13 | Lonicera_rupicola36511 | 31.949722 | 98.595 | 3720 | Hengduan | 36511 |
| 14 | Lonicera_rupicola32079 | 31.092778 | 96.419444 | 4400-4620 | Hengduan | 32079 |
| 15 | Lonicera_rupicola202113 | 30.97405 | 79.03297 | 3668 | WII | WII20513 |
| 16 | Lonicera_rupicola292 | 34.3575 | 100.255556 | 3870 | MO | 4647484 |
| 17 | Lonicera_syringantha130 | 32.598147 | 102.445437 | 3570 | K | 1988-8493 |
| 18 | Lonicera_syringantha177 | 34.648889 | 100.610556 | 3500-3600 | MO | 4648115 |
| 19 | Lonicera_rupicola362 | 32.95 | 103.713297 | 3300 | MO | MW0748589 |
| 20 | Lonicera_rupicola88 | 34.494444 | 101.169167 | 3410 | E | E00137326 |
| 21 | Lonicera_rupicola156 | 32.343333 | 100.768889 | 3570 | F | 2308854 |
| 22 | Lonicera_rupicola324 | 34.059444 | 102.655556 | 3570 | F | 2308850 |
| 23 | Lonicera_rupicola322 | 32.233889 | 101.219722 | 3760 | F | 2308842 |
| 24 | Lonicera_rupicola157 | 33.421391 | 104.396539 | 3658 | MO | 4665931 |
| 25 | Lonicera_syringantha313 | 32.0475 | 102.016944 | 3535 | F | 2308855 |
| 26 | Lonicera_syringanthaN1412 | 27.8789386 | 86.81432362 | 4149 | KATH | N14; No<br>voucher |
| 27 | Lonicera_syringantha199 | 36.766667 | 102.533333 | 2735 | E | E00137329 |
| 28 | Lonicera_minuta273 | 33.693889 | 99.431111 | 4050 | E | E01059975 |
| 29 | Lonicera_rupicola174 | 32.066667 | 97.05 | 3910 | MO | zero5093630 |
| 30 | Lonicera_syringanthaSL2 | 28.02871204 | 88.70727423 | 3690 | WII | WII20504 |
| 31 | Lonicera_rupicola323 | 33.3025 | 101.527222 | 3720 | F | 2308284 |
| 32 | Lonicera_rupicola181 | 30.961111 | 98.306944 | 3600 | YU | 41497 |
| 33 | Lonicera_syringantha293 | 35.6778 | 102.911 | 3658 | MO | 4666022 |
| 34 | Lonicera_syringantha48 | 33.108611 | 102.631944 | 3485-3520 | F | 2308830 |
| 35 | Lonicera_rupicola311 | 28.383333 | 99.016667 | 4050-4100 | F | 2279693 |
| 36 | Lonicera_minuta318 | 35.261667 | 98.9825 | 4250 | MO | 4648043 |
| 37 | Lonicera_syringantha363 | 32.59231 | 103.38686 | 3962 | MW | MW0748590 |
| 38 | Lonicera_rupicola331 | 28.3372502 | 99.0870126 | 4236 | KUN | KUN 1492702 |
| 39 | Lonicera_rupicola310 | **34.93917** | 103.73472 | 2880 | MO | 4666088 |
| 40 | Lonicera_rupicola309 | 30.040278 | 101.838333 | 3700-3750 | MO | 5768591 |
| 41 | Lonicera_syringantha175 | 35.114999 | 101.401429 | None | MO | 4560870 |
| 42 | Lonicera_rupicola301 | 31.878056 | 99.763889 | 3690 | F | 2281219 |
| 43 | Lonicera_minuta105 | 34.516111 | 100.220556 | 3675 | E | E00137324 |
| 44 | Lonicera_syringantha321 | 32.718056 | 101.937222 | 3175 | F | 2308853 |
| 45 | Lonicera_rupicola277 | 32.988278 | 100.702556 | 3266 | KUN | 0714392 |
| 46 | Lonicwera_myrtillus319 | 29.166667 | 83.95 | 3950 | MO | 6154886 |
| 47 | Lonicera_angustifolia1 | 27.516675 | 88.183782 | 3760 | WII | WII21209 |
| 48 | Lonicera_tubuliflora | 27.928733 | 88.732885 | 4238 | WII | WII21207 |
| 49 | Lonicera_angustifoliaSurk | 30.41009 | 78.28813 | 2640 | WII | WII20518 |
| 50 | Lonicera_angustifolia2019T5 | 27.50735 | 88.17826 | 2980 | WII | WII20519 |
| 51 | Lonicera_angustifolia2019Roh | 32.45042 | 77.14383 | 2900 | WII | Roh; no<br>voucher |
| 52 | Lonicera_myrtillus14 | 30.9914 | 78.99503 | 3304 | WII | WII20502 |
| 53 | Lonicera_angustifolia202119b | 30.76873 | 79.80279 | 3680 | WII | WII20527 |
| 54 | Lonicera_myrtillus202138 | 27.514098 | 92.094103 | 4170 | WII | WII20526 |
| 55 | Lonicera_myrtillus20213 | 33.99901 | 75.38319 | 3276 | WII | 3; no voucher |
| 56 | Lonicera_myrtillus202112 | 30.96901 | 79.03784 | 3721 | WII | 12; no voucher |
| 57 | Lonicera angustifolia201918 | 30.62924 | 78.93575 | 3140 | WII | 18; no voucher |
| 58 | Lonicera angustifolia20219 | 30.747493 | 77.850481 | 3200 | WII | WII20512 |
| 59 | Lonicera_myrtillus135 | 30.80694 | 79.80777164 | 3752 | WII | WII20528 |
| 60 | Lonicera_myrtilloides350 | 30.249728 | 80.529964 | 3300 | WII | 350; no<br>voucher |
| 61 | *Lonicera angustifolia*270 | 28.601389 | 84.635 | 3540 | E | E00625377 |
| 62 | *Lonicera myrtillus*141Sur | 33.143814 | 76.42034149 | 2900 | WII | Sur; no<br>voucher |
| 63 | *Lonicera_myrtilloides*N4370 | 28.6192538 | 84.14005798 | 3195 | KATH | KATH141148 |
| 64 | *Lonicera_angustifolia*230 | 28.631389 | 84.637222 | 3690 | E | E00654717 |
| 65 | *Lonicera_angustifolia*N76 | 28.21247735 | 85.490246 | 3241 | KATH | KATH141146 |
| 66 | *Lonicera_myrtillus*144 | 27.78379585 | 96.88418615 | 3393 | WII | WII20509 |
| 67 | *Lonicera_angustifolia*127 | None | None | None | K | 1989-8329 |
| 68 | *Lonicera_angustifolia*N109 | 29.26104489 | 82.10572724 | 2918 | KATH | KATH140530 |
| 69 | *Lonicera_angustifolia*128 | None | None | None | K | 171/K0005687<br>14 |
| 70 | *Lonicera_myrtillus*N1211 | 27.88064445 | 86.81667377 | 4154 | KATH | KATH141142 |
| 71 | *Lonicera_myrtillus*N98 | 28.21277654 | 85.53123363 | 3543 | KATH | KATH141144 |
| 72 | *Lonicera_myrtillus*152 | 29.809385 | 95.7466 | None | KUN | KUN 1521957 |
| 73 | *Lonicera_angustifolia*N1110 | 27.80762337 | 86.71972431 | 3513 | KATH | KATH141140 |
| 74 | *Lonicera_myrtillus*134 | 27.486723 | 90.902638 | None | E | 19891047 C |
| 75 | *Lonicera_myrtillus*196 | 29.066667 | 83.866667 | 4030 | E | E00228930 |
| 76 | *Lonicera_myrtillus*N2368 | 28.66367288 | 84.0199848 | 3488 | KATH | KATH141136 |
| 77 | *Lonicera angustifolia*201915 | 30.45692 | 78.35290 | 3140 | WII | 18; no voucher |
| 78 | *Lonicera_spinosa*20217a | 33.7359 | 77.74339 | 3740 | WII | 7a; no voucher |
| 79 | *Lonicera_spinosa*201911 | 32.78408 | 77.35111 | 4200 | WII | WII20524 |
| 80 | *Lonicera_spinosa*202116 | 32.10406 | 78.18567 | 3388 | WII | WII20521 |
| 81 | *Lonicera_spinosa*20217g | 33.26757 | 78.47359 | 3969 | WII | 7g; no voucher |
| 82 | *Lonicera_spinosa*20217f | 32.96373 | 78.25736 | 4525 | WII | 7f; no voucher |
| 83 | *Lonicera_spinosa*20217e | 33.19333333 | 78.4641667 | 4600 | WII | 7e; no voucher |
| 84 | Lonicera_spinosa20216b | 34.39643 | 77.66162 | 3990 | WII | 6b; no voucher |
| 85 | Lonicera_spinosa20215a | 34.158728 | 76.842328 | 3809 | WII | 5a; no voucher |
| 86 | Lonicera_spinosa20215c | 34.070472 | 76.828 | 3857 | WII | 5c; no voucher |
| 87 | Lonicera_alberti202117 | 32.341964 | 77.924421 | 4340 | WII | WII20522 |
| 88 | Lonicera_spinosa20217c | 33.33111111 | 78.45916667 | 4670 | WII | 7c; no voucher |
| 89 | Lonicera_spinosa202118 | 32.439751 | 77.785971 | 4,090 | WII | WII20523 |
| 90 | Lonicera_spinosa20214 | 34.33157 | 76.57642 | 3540 | WII | 4; no voucher |
| 91 | Lonicera_spinosa20217i | 33.39994 | 78.24206 | 4010 | WII | 7i; no voucher |
| 92 | Lonicera_spinosa20217d | 32.99465 | 78.26197 | 4442 | WII | 7d; no voucher |
| 93 | Lonicera_spinosa20216a | 34.3547 | 77.66444 | 4206 | WII | 6a; no voucher |
| 94 | Lonicera_spinosa20217h | 33.25929 | 78.49787 | 4166 | WII | 7h; no voucher |
| 95 | Lonicera_spinosa20217b | 33.66788 | 77.73195 | 3990 | WII | 7b; no voucher |
| 96 | Lonicera_spinosa1 | 32.79256 | 77.32874 | 4200 | WII | WII20524 |
| 97 | Lonicera_alberti150 | 42.866458 | 80.499236 | 723 | KUN | 1462668 |
| 98 | Lonicera_tsarongensis1 | 27.800995 | 88.770825 | 3063 | WII | WII21206 |
| 99 | Lonicera_tsarongensisN6371 | 28.21256706 | 85.48981776 | 3233 | KATH | KATH140535 |
| 100 | Lonicera_tomentella133 | 28 | 89.783333 | None | E | 19841817 |
| 101 | Lonicera_tomentellaSL113 | 27.79635747 | 88.5529237 | 3168 | WII | WII20503 |
| 102 | Lonicera_tomentellaSL414 | 27.71279367 | 88.5557661 | 2608 | WII | WII20506 |
| 103 | Lonicera_tsarongensisN8372 | 28.2136461 | 85.52998855 | 3533 | KATH | KATH140533 |
| 104 | Lonicera_arborea | none | none | none | MA | BEEE46 |
| 105 | Lonicera_oblongifolia | none | none | none | YU | no voucher |

## APPENDIX S1. Morphological characters analyzed

The characters and their states are as follows:

1. ***Floral symmetry***: (0) bilaterally symmetrical, (1) radially symmetrical. We scored species having the 4:1 flower form, one to three nectaries and/or corolla tubes with one-sided gibbosity as bilateral. All *Isoxylosteum* species have radially symmetrical flowers with five nectaries. Some *L. angustifolia* flowers have a slightly curved corolla tube (Fig. 6A), but these were scored as radial based on their predominantly uncurved corolla tubes, the presence of five nectaries, and the absence of one-sided gibbosity.
2. ***Thorns:*** (0) absent, (1) present.
3. ***Number of leaves per node***: (0) two, leaves opposite, (1) three, leaves whorled.
4. ***Lower (abaxial) leaf surface:*** (0) glabrous (Fig. 6P), (1) lanate (Fig. 6R).
5. ***Secondary leaf veins conspicuous:*** (0) yes (Figs. 6P,Q,R), (1) no (Figs. 6S,T).
6. ***Bracteoles fusion:*** We followed the scoring of Srivastav et al. (2023): (0) Bracteoles of the lateral flowers present and free from one another, (1) bracteoles absent or highly reduced, (2) bracteoles of the same flower fused, (3) bracteoles of adjacent flowers fused in pairs, (4) all bracteoles fused into a cup that partially covers the pair of adjacent ovaries (partial cupule; Figs. 6A-E), and (5) all bracteoles fused into a cupule that completely envelops the pair of adjacent ovaries (i.e., a complete cupule).
7. ***Ovary fusion:*** We followed the scoring of Srivastav et al. (2023): (0) Ovaries free or fused only at the base (less than one-quarter of the mature ovary length; Figs. 6 M-O),
8. ovaries partially fused (between one-quarter and three-quarters of the mature ovary length), (2) ovaries fully fused (more than three-quarters of the ovary length or fully fused; Figs. 6 K,L).
9. ***Corolla lobe number:*** (0) five, (1) four to seven.
10. ***Pubescence inside corolla tube:*** (0) present, (1) absent. Species with trichomes visible on the inner surface of the corolla were scored as being pubescent. Those with no trichomes or trichomes present only at the base of the filaments and style were scored as lacking pubescence.
11. ***Stigma-anther position:*** (0) stigma and anthers exserted (Fig. 6I), (1) stigma and anthers deeply seated inside the corolla tube and hidden from view (Figs. 6F, H), (2) stigma and anthers reaching the opening of the corolla tube such that they are both visible (Fig. 6G), (3) stigma deeply seated inside of the corolla tube and anthers reaching the opening of the corolla tube (Fig. 6J).
12. ***Fruit color:*** (0) red (Figs. 6 K,M), (1) blue with a waxy bloom (Fig. 6L), (2) translucent purple, (3) black, (4) white, (5) yellow or yellowish-orange (Fig. 6N).

## APPENDIX S2. Key to the species of *Isoxylosteum*

Our findings on character evolution can readily be translated into a taxonomic key to distinguish the five species of *Isoxylosteum* recognized here:

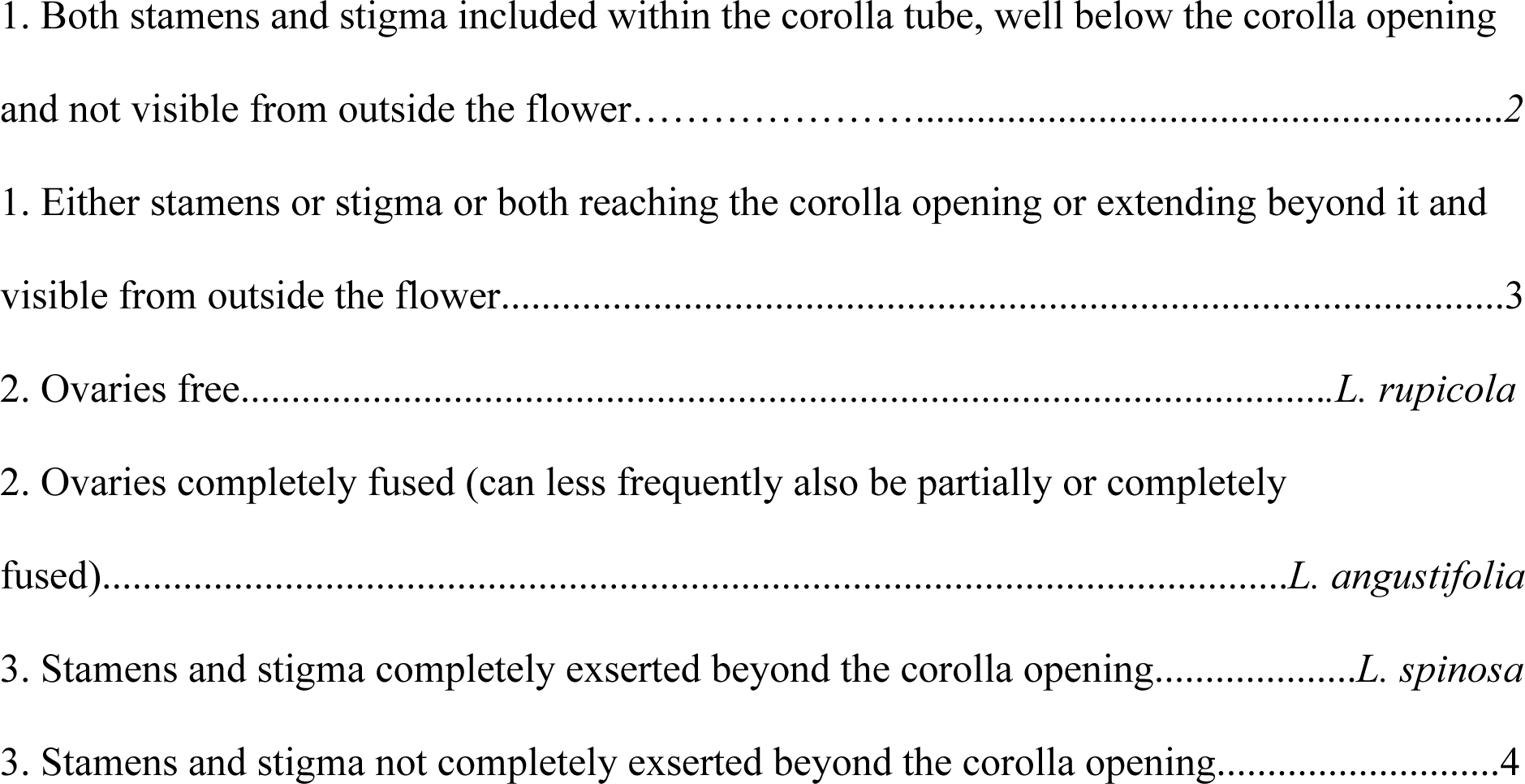
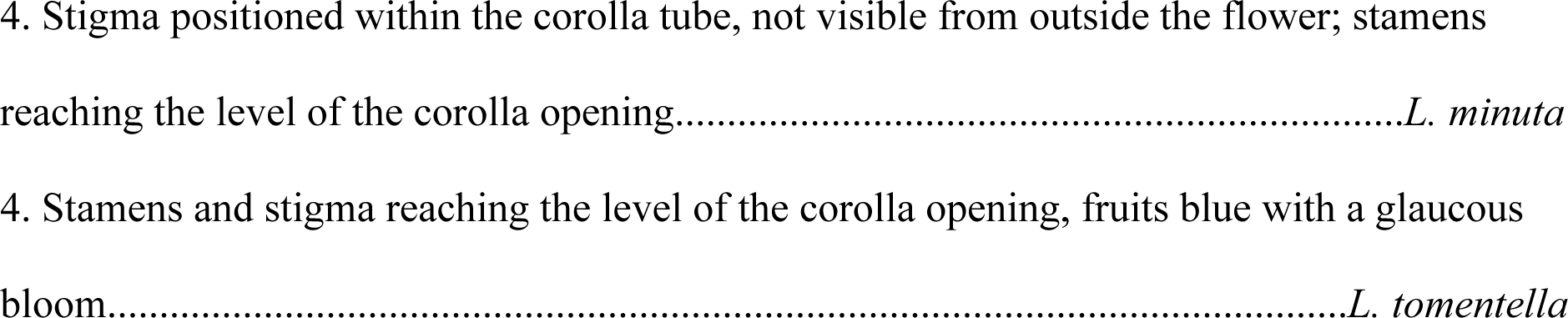

**TABLE S1.**
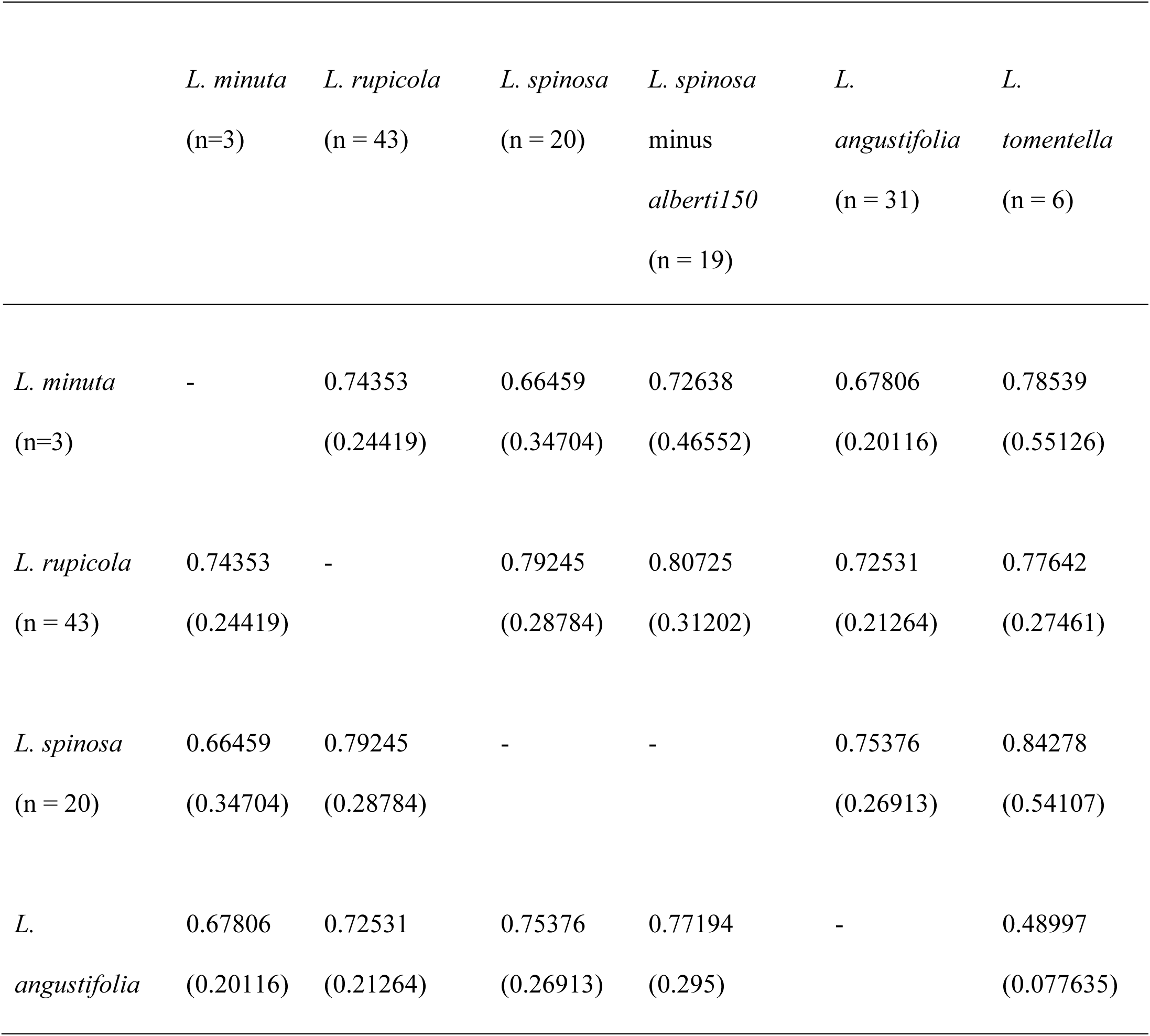
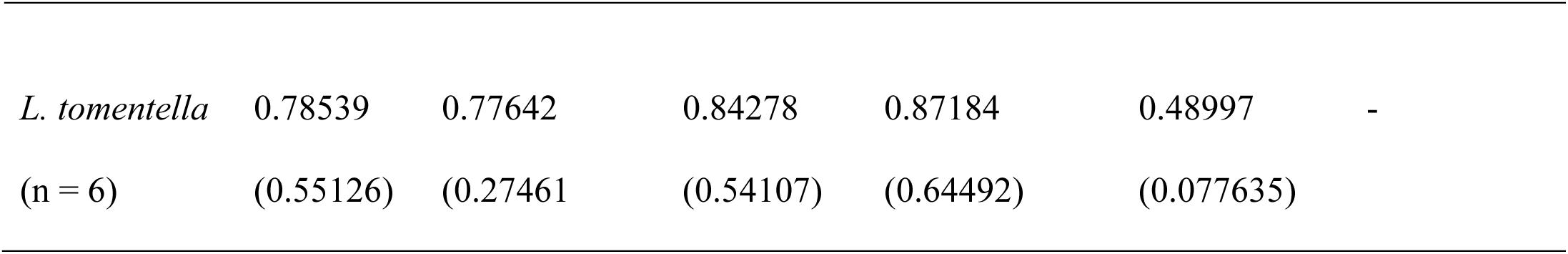
Weighted F_ST_ values between the five delimited species of *Isoxylosteum* with and without the *L. spinosa* (alberti150) accession from Central Asia. In parenthesis are mean F_ST_ values.

|  | *L. minuta*<br>(n=3) | *L. rupicola*<br>(n = 43) | *L. spinosa*<br>(n = 20) | *L. spinosa*<br>minus<br>*alberti150*<br>(n = 19) | *L.*<br>*angustifolia*<br>(n = 31) | *L.*<br>*tomentella*<br>(n = 6) |
| --- | --- | --- | --- | --- | --- | --- |
| *L. minuta*<br>(n=3) | - | 0.74353<br>(0.24419) | 0.66459<br>(0.34704) | 0.72638<br>(0.46552) | 0.67806<br>(0.20116) | 0.78539<br>(0.55126) |
| *L. rupicola*<br>(n = 43) | 0.74353<br>(0.24419) | - | 0.79245<br>(0.28784) | 0.80725<br>(0.31202) | 0.72531<br>(0.21264) | 0.77642<br>(0.27461) |
| *L. spinosa*<br>(n = 20) | 0.66459<br>(0.34704) | 0.79245<br>(0.28784) | - | - | 0.75376<br>(0.26913) | 0.84278<br>(0.54107) |
| *L.*<br>*angustifolia* | 0.67806<br>(0.20116) | 0.72531<br>(0.21264) | 0.75376<br>(0.26913) | 0.77194<br>(0.295) | - | 0.48997<br>(0.077635) |
| *L. tomentella* | 0.78539 | 0.77642 | 0.84278 | 0.87184 | 0.48997 | - |
| (n = 6) | (0.55126) | (0.27461 | (0.54107) | (0.64492) | (0.077635) |  |

**TABLE S2.** Results of Welch’s one-way ANOVA testing for differences in leaf traits among five *Isoxylosteum* species.

| Leaf trait | Welch's F | df<sub>1</sub> | df<sub>2</sub> | p-value |
| --- | --- | --- | --- | --- |
| Leaf length | 14.528 | 4 | 41.617 | <0.001 |
| Leaf width | 72.595 | 4 | 41.944 | <0.001 |
| Leaf length-to-width ratio | 51.175 | 4 | 34.597 | <0.001 |

**TABLE S3.** Summary of significant differences in leaf traits among species. Values indicate the number of significant species-pair comparisons out of 10 possible pairwise comparisons, based on Games–Howell post hoc tests following Welch’s one-way ANOVA.

| Leaf trait | No. significant pairwise differences (out of 10) |
| --- | --- |
| Leaf length | 6 |
| Leaf width | 8 |
| Leaf length-to-width ratio | 8 |

**TABLE S4.** LDA leave-one-out cross-validation confusion matrix based on leaf traits.

| Actual species | *L. angustifolia* | *L. tomentella* | *L. rupicola* | *L. spinosa* | *L. minuta* | Total | Correct (%) |
| --- | --- | --- | --- | --- | --- | --- | --- |
| *L. angustifolia* | 40 | 5 | 4 | 2 | 0 | 51 | 78.4 |
| *L. tomentella* | 9 | 13 | 0 | 0 | 0 | 22 | 59.1 |
| *L. rupicola* | 23 | 2 | 8 | 3 | 0 | 36 | 22.2 |
| *L. spinosa* | 2 | 0 | 1 | 21 | 0 | 24 | 87.5 |
| *L. minuta* | 3 | 0 | 1 | 3 | 0 | 7 | 0.0 |

**TABLE S5.** Loadings of floral traits on the first four principal components.

| Floral trait | PC1 | PC2 | PC3 | PC4 |
| --- | --- | --- | --- | --- |
| Corolla length | 0.396 | 0.154 | 0.192 | 0.031 |
| Corolla tube length | 0.374 | 0.043 | 0.270 | 0.000 |
| Corolla tube width at midpoint | -0.108 | -0.485 | -0.054 | -0.087 |
| Corolla lobe length | 0.389 | 0.185 | 0.087 | 0.037 |
| Corolla lobe width | 0.327 | -0.068 | 0.083 | 0.103 |
| Corolla diameter | 0.393 | 0.096 | 0.124 | 0.076 |
| Peduncle length | 0.002 | -0.179 | 0.530 | -0.588 |
| Calyx tube length | 0.306 | -0.032 | -0.340 | 0.019 |
| Calyx lobe length | 0.176 | -0.024 | -0.533 | -0.564 |
| Calyx lobe width | 0.244 | -0.012 | -0.323 | 0.126 |
| Ovary length | 0.255 | -0.375 | 0.026 | -0.334 |
| Ovary breadth | 0.042 | -0.532 | 0.192 | 0.376 |
| Calyx tube width at base | 0.169 | -0.484 | -0.185 | 0.200 |

**TABLE S6.** LDA results based on floral traits.

| Actual species | *L. angustifolia* | *L. rupicola* | *L. spinosa* | *L. tomentella* | Total | Correct (%) |
| --- | --- | --- | --- | --- | --- | --- |
| *L. angustifolia* | 51 | 1 | 0 | 1 | 53 | 96.2 |
| *L. rupicola* | 1 | 33 | 1 | 1 | 36 | 91.7 |
| *L. spinosa* | 1 | 2 | 22 | 0 | 25 | 88.0 |
| *L. tomentella* | 6 | 0 | 1 | 15 | 22 | 68.2 |

B. LDA coefficients.
| Floral PC | LD1 | LD2 | LD3 |
| --- | --- | --- | --- |
| PC1 | 0.633 | -0.393 | -0.308 |
| PC2 | 0.418 | -0.578 | 0.702 |
| PC3 | -1.155 | -0.697 | -0.050 |
| PC4 | -0.784 | -0.815 | -0.312 |
| Proportion of trace | 0.606 | 0.293 | 0.102 |

**TABLE S7.**
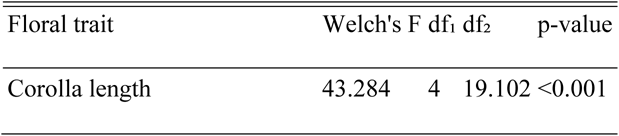
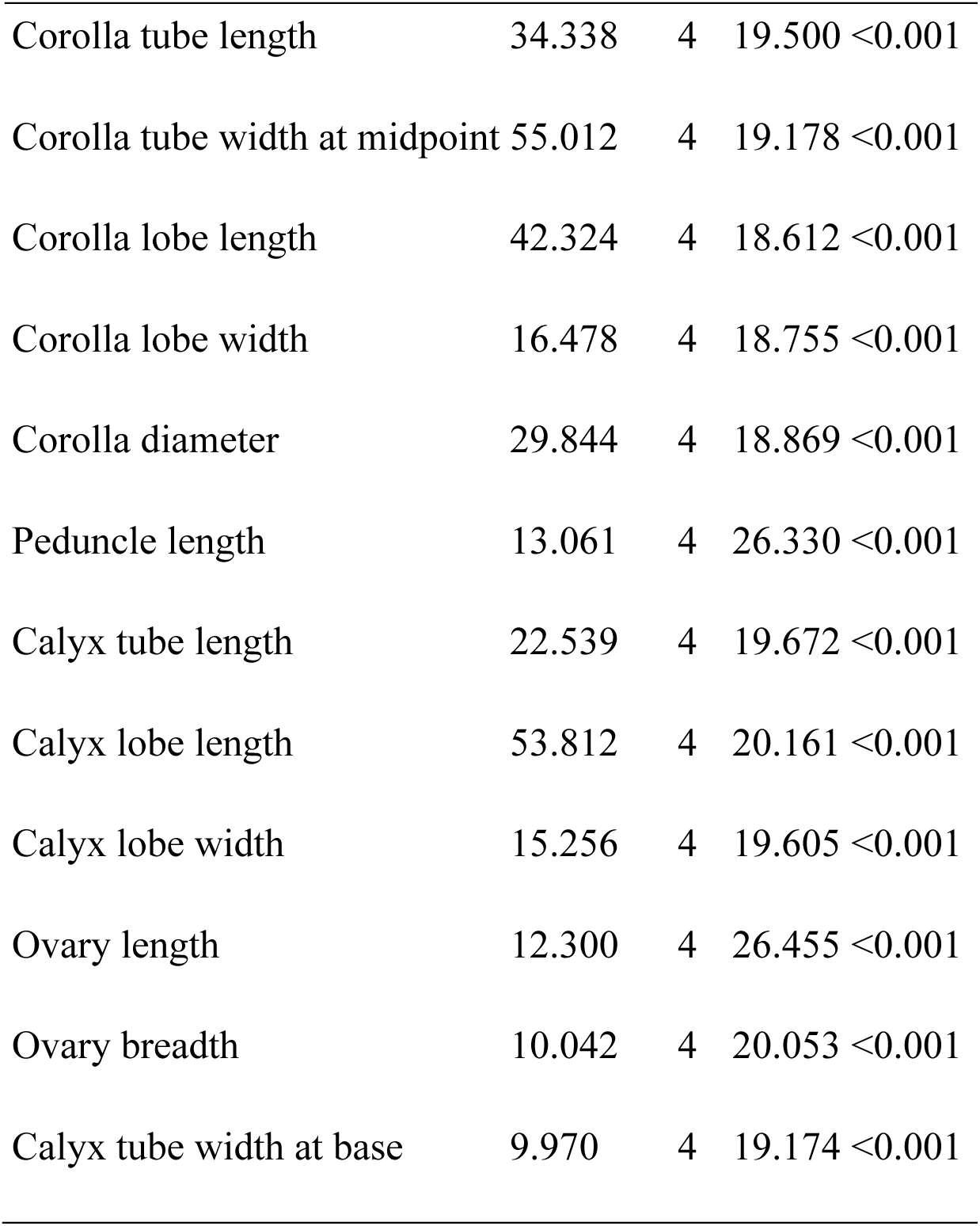
Results of Welch’s one-way ANOVA testing for differences in floral traits among five *Lonicera* species.

**TABLE S8.** Summary of significant differences in floral traits among species.

| Floral trait | No. significant pairwise differences (out of 10) |
| --- | --- |
| Corolla length | 5 |
| Corolla tube length | 5 |
| Corolla tube width at midpoint | 4 |
| Corolla lobe length | 5 |
| Corolla lobe width | 3 |
| Corolla diameter | 4 |
| Peduncle length | 6 |
| Calyx tube length | 3 |
| Calyx lobe length | 8 |
| Calyx lobe width | 3 |
| Ovary length | 4 |
| Ovary breadth | 5 |
| Calyx tube width at base | 4 |

| Species comparison | No. significant pairwise differences (out of 13) |
| --- | --- |
| *L. angustifolia* vs. *L. minuta* | 3 |
| *L. angustifolia* vs. *L. rupicola* | 12 |
| *L. angustifolia* vs. *L. spinosa* | 10 |
| *L. angustifolia* vs. *L. tomentella* | 10 |
| *L. minuta* vs. *L. rupicola* | 1 |
| *L. minuta* vs. *L. spinosa* | 1 |
| *L. minuta* vs. *L. tomentella* | 3 |
| *L. rupicola* vs. *L. spinosa* | 8 |
| *L. rupicola* vs. *L. tomentella* | 4 |
| *L. spinosa* vs. *L. tomentella* | 7 |

**TABLE S9.** Games–Howell post hoc comparisons of floral traits between ***L. tubuliflora*** and *L. angustifolia* and *L. rupicola*. Values are adjusted *P*-values; NS = not

| Floral trait | *L. tubuliflora* vs. *L. angustifolia* | *L. tubuliflora* vs. *L. rupicola* |
| --- | --- | --- |
| Corolla length | 0.0023 | NS |
| Corolla tube length | 0.0177 | NS |
| Corolla tube width at midpoint | NS | NS |
| Corolla lobe length | NS | 0.0183 |
| Corolla lobe width | NS | NS |
| Corolla diameter | NS | 0.0461 |
| Peduncle length | NS | NS |
| Calyx tube length | NS | 0.0135 |
| Calyx lobe length | NS | <0.001 |
| Calyx lobe width | NS | NS |
| Ovary length | NS | NS |
| Ovary breadth | NS | NS |
| Calyx tube width at base | NS | NS |

**TABLE S10.**
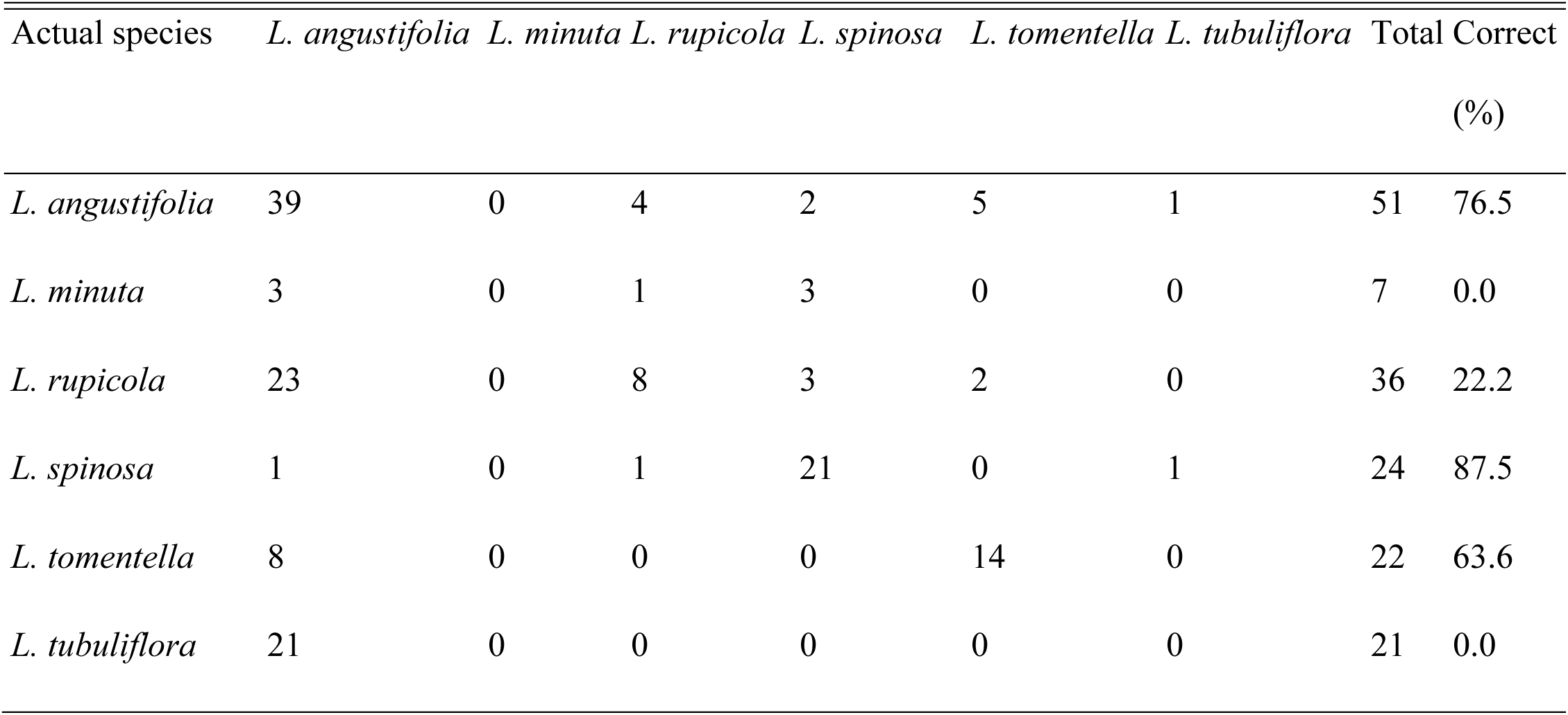
LDA leave-one-out cross-validation confusion matrix of leaf traits with *L. tubuliflora* included.

**TABLE S11.** Number of significant leaf trait differences between species pairs Values indicate the number of leaf traits showing significant differences between each species pair out of 2 traits (leaf length and width), based on Games–Howell post hoc tests following Welch’s one-way ANOVA.

| Species comparison | No. significant pairwise differences (out of 2) |
| --- | --- |
| *L. angustifolia* vs. *L. minuta* | 2 |
| *L. angustifolia* vs. *L. rupicola* | 0 |
| *L. angustifolia* vs. *L. spinosa* | 1 |
| *L. angustifolia* vs. *L. tomentella* | 2 |
| *L. minuta* vs. *L. rupicola* | 2 |
| *L. minuta* vs. *L. spinosa* | 0 |
| *L. minuta* vs. *L. tomentella* | 2 |
| *L. rupicola* vs. *L. spinosa* | 1 |
| *L. rupicola* vs. *L. tomentella* | 2 |
| *L. spinosa* vs. *L. tomentella* | 2 |

**FIGURE S1.**
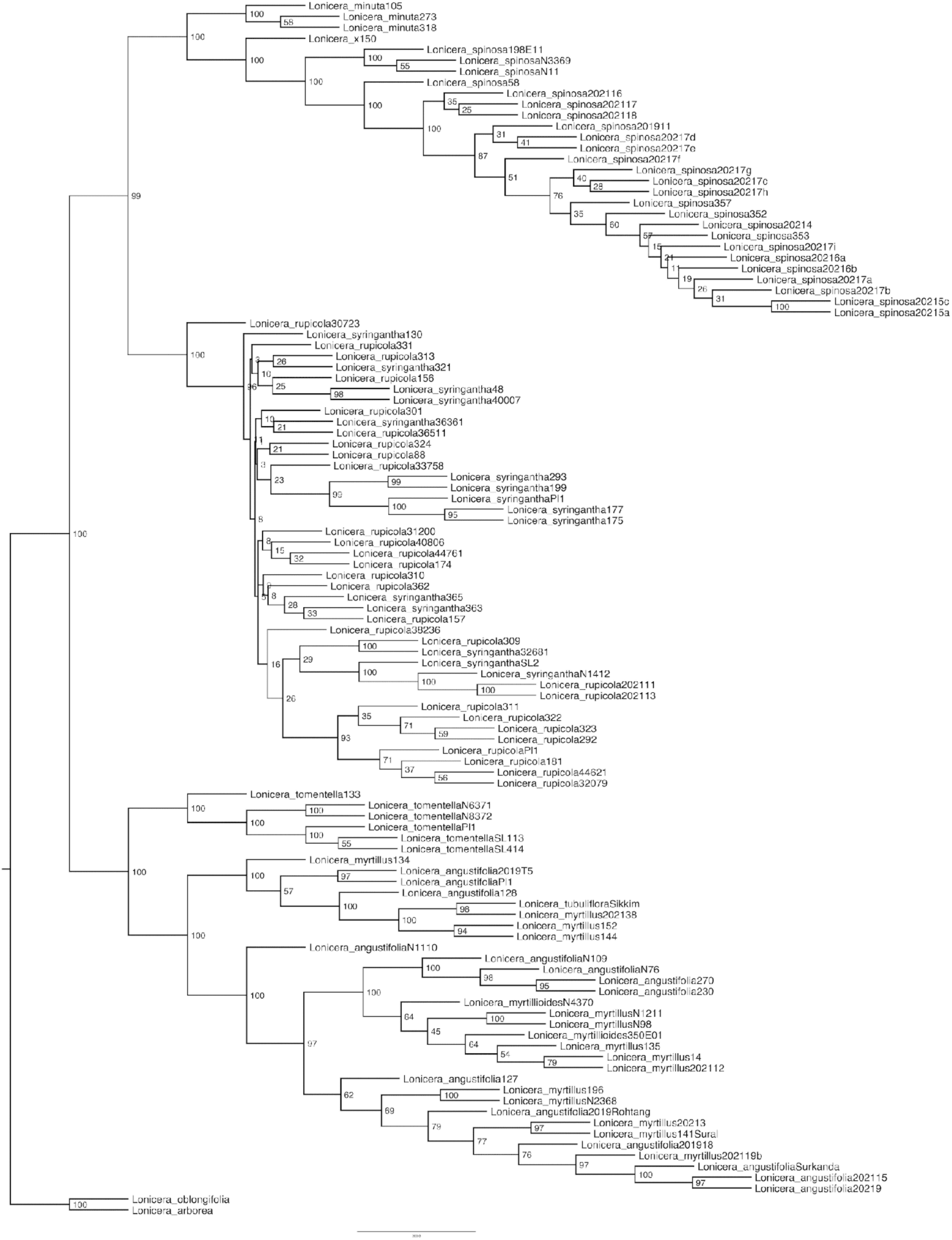
Tetrad tree of *Isxoylosteum* built using m4 assembly.

**FIGURE S2.**
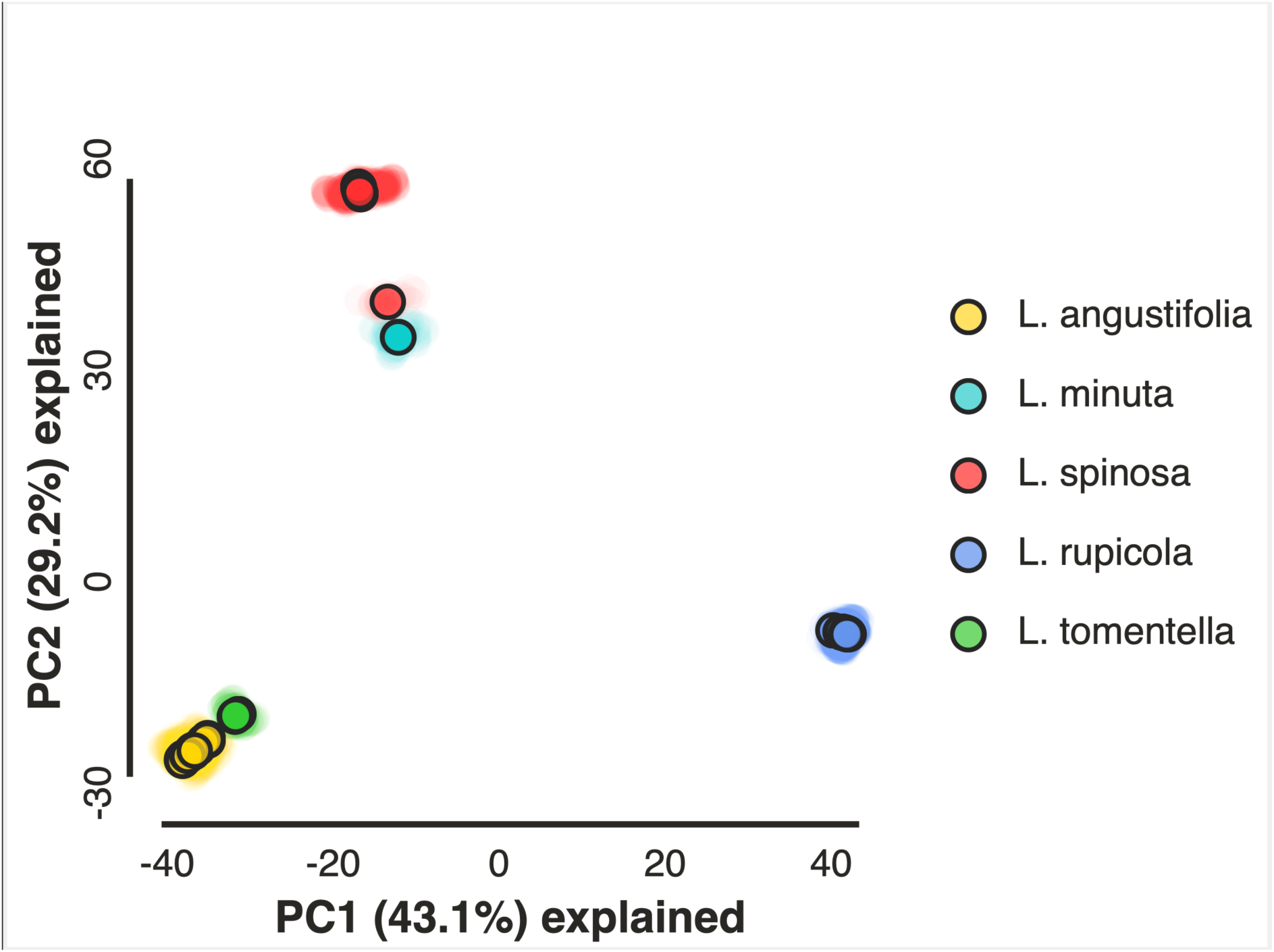
Principal component analysis (PCA) of genetic variance for *Isoxylosteum* using the sample method where populations were assigned to the five major clades.

**FIGURE S3.**
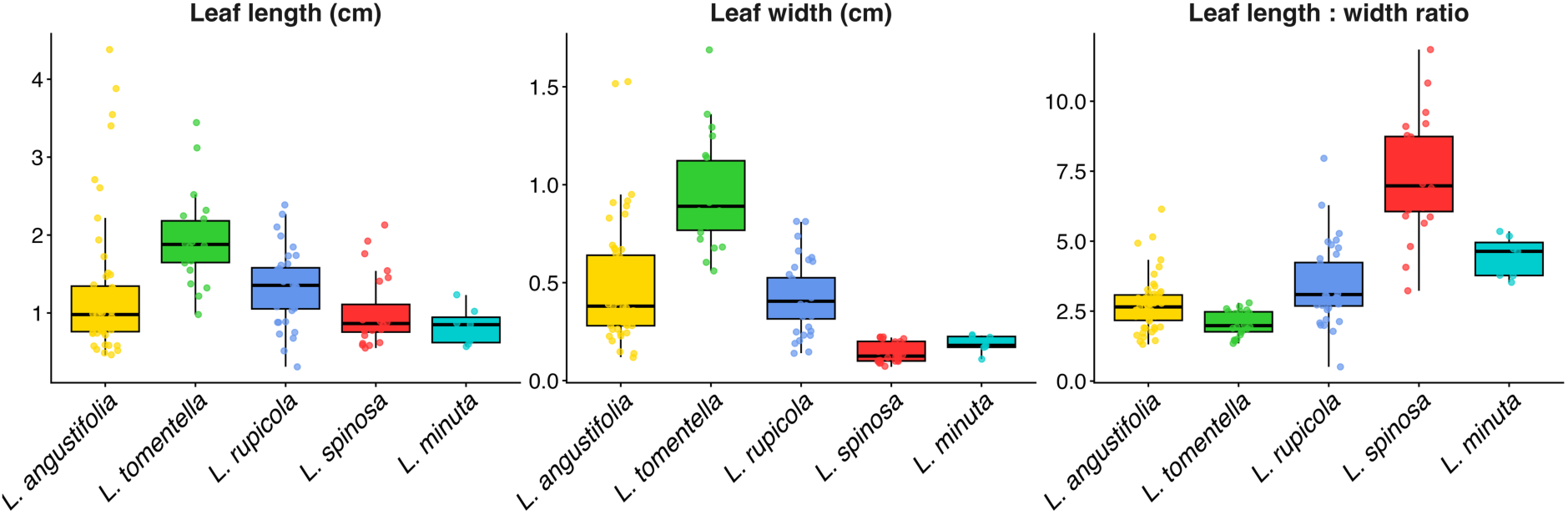
Variation in leaf length, width, and length-to-width ratio among *Isoxylosteum* species.

**FIGURE S4.**
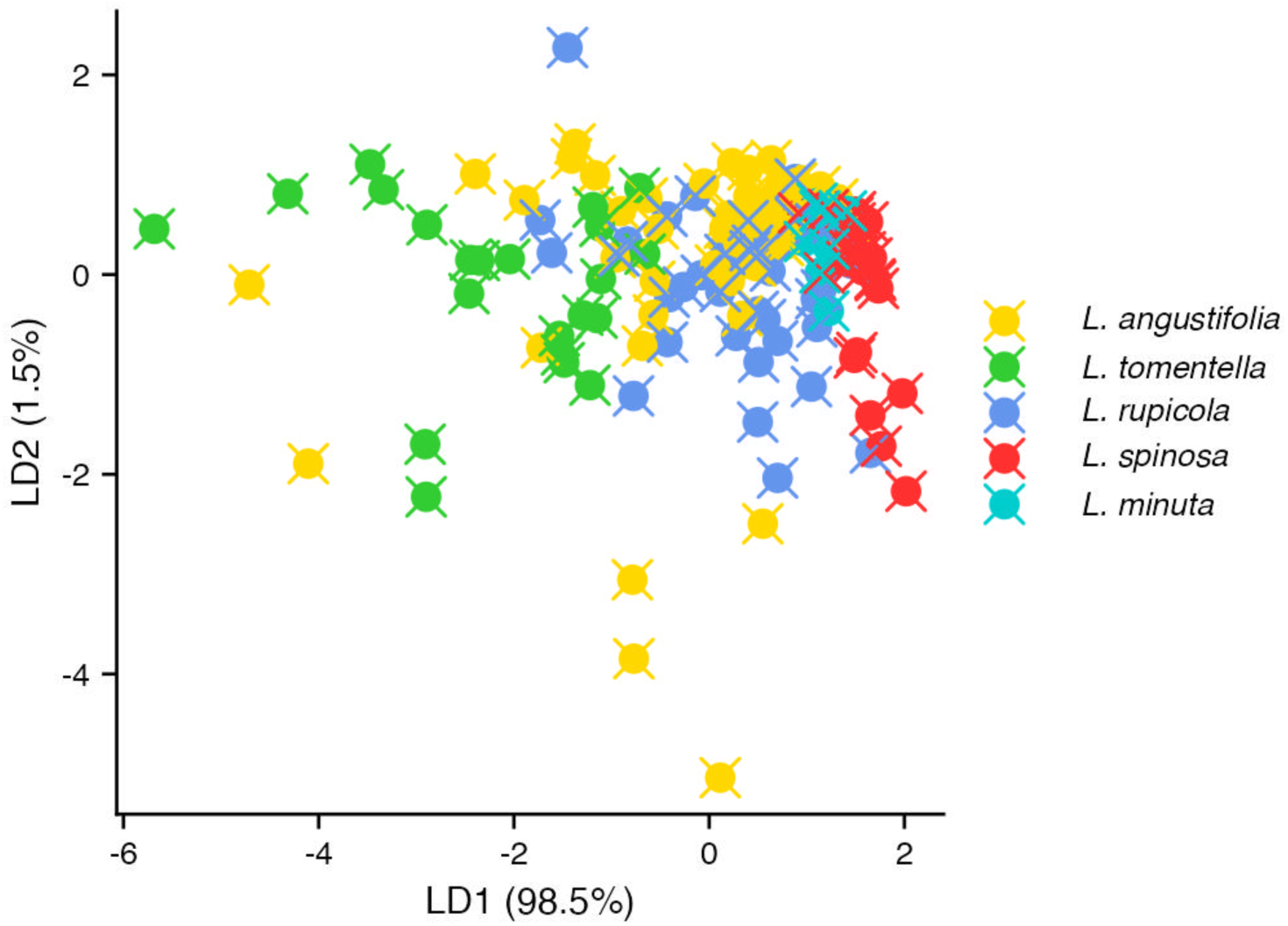
LDA plot for leaf length and width.

**FIGURE S5.**
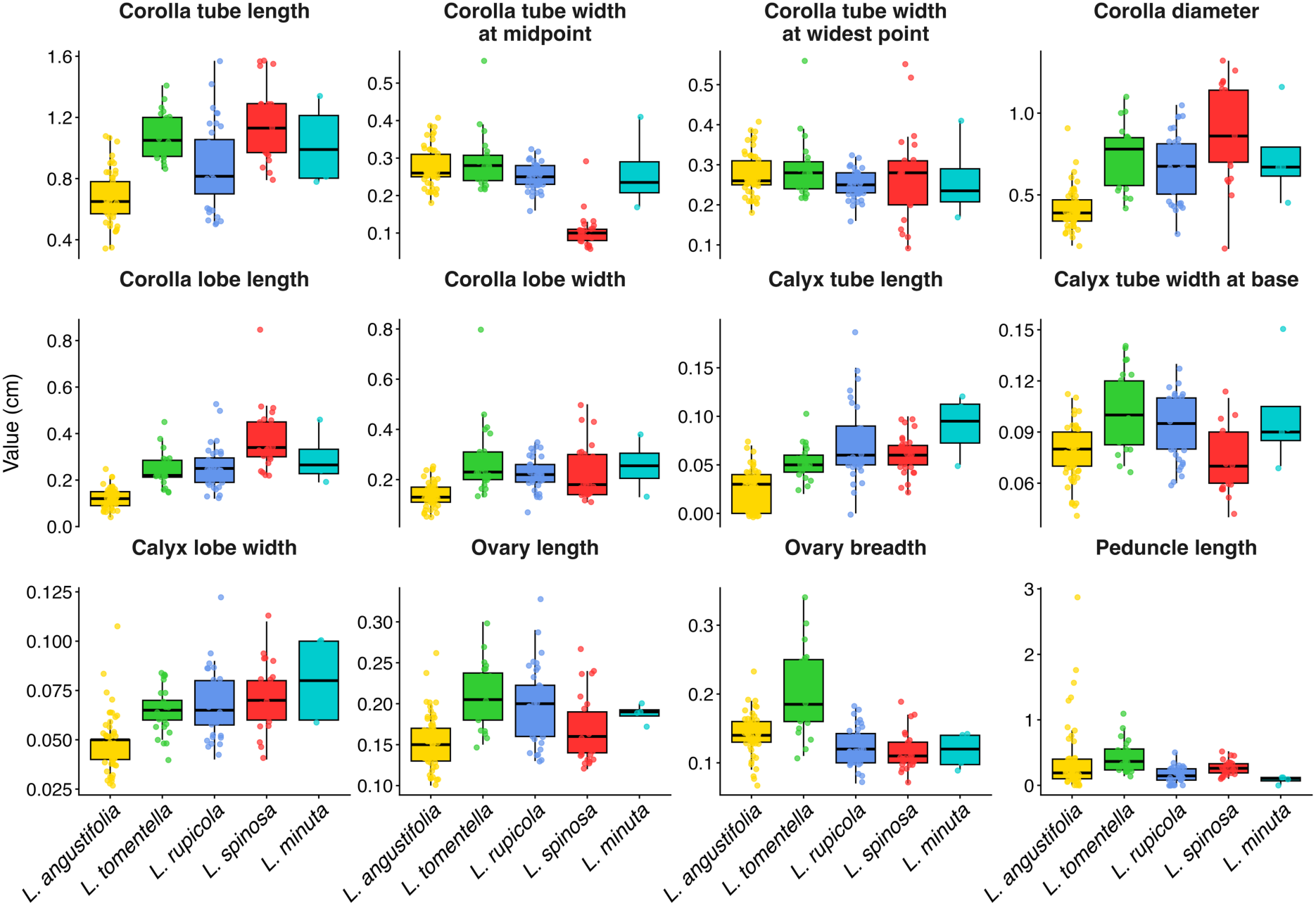
Boxplots showing variation in floral traits.

**FIGURE S6.**
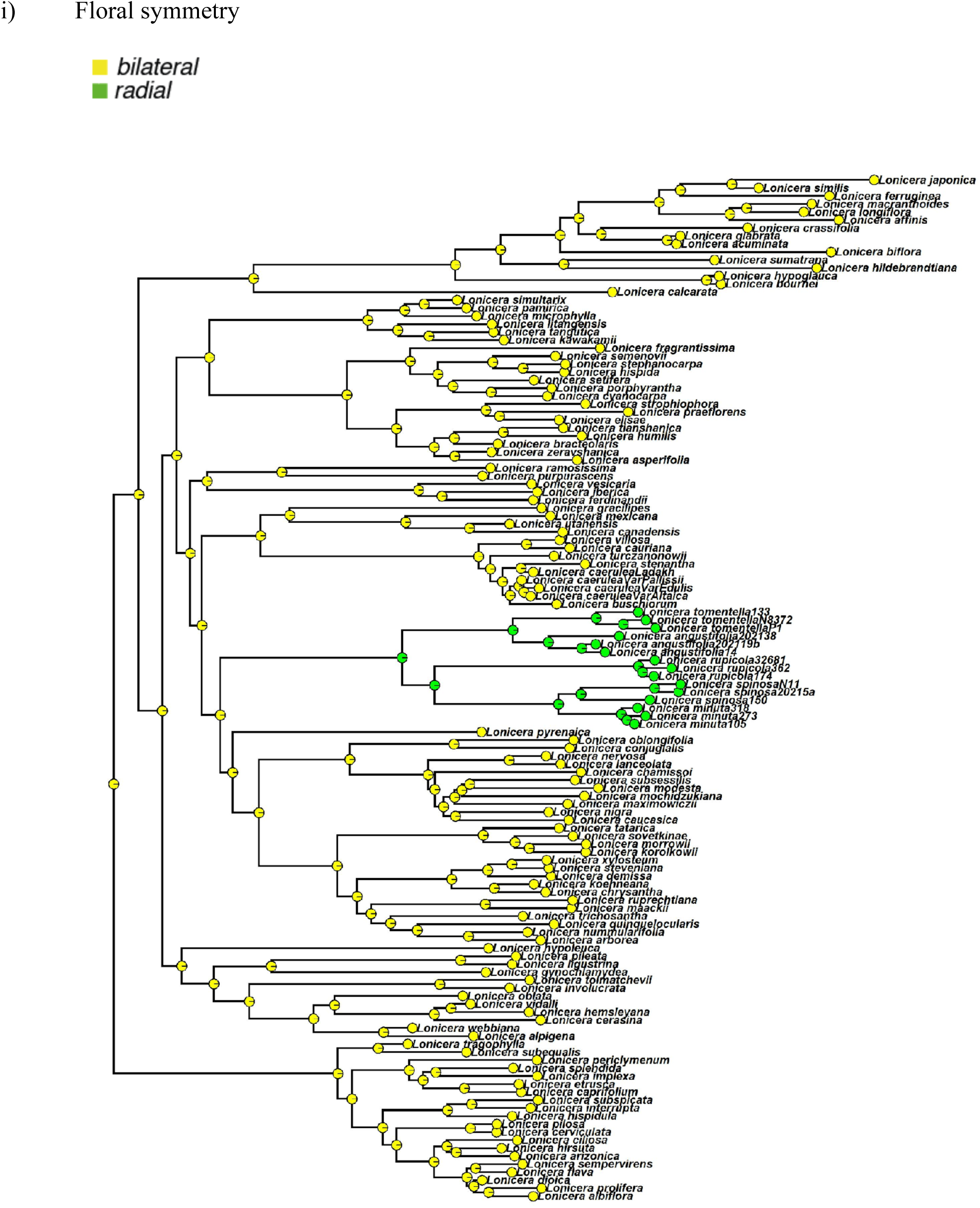
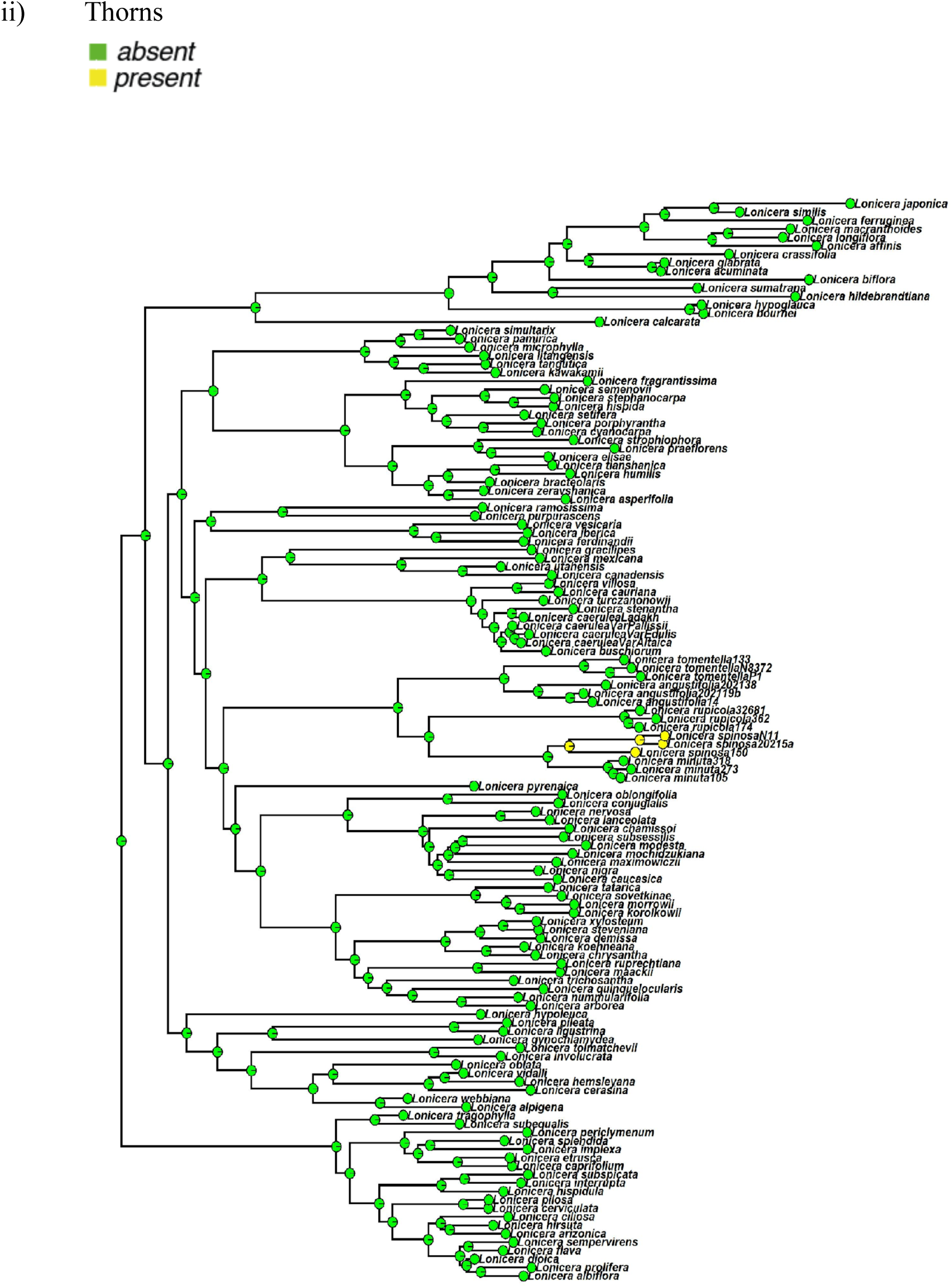
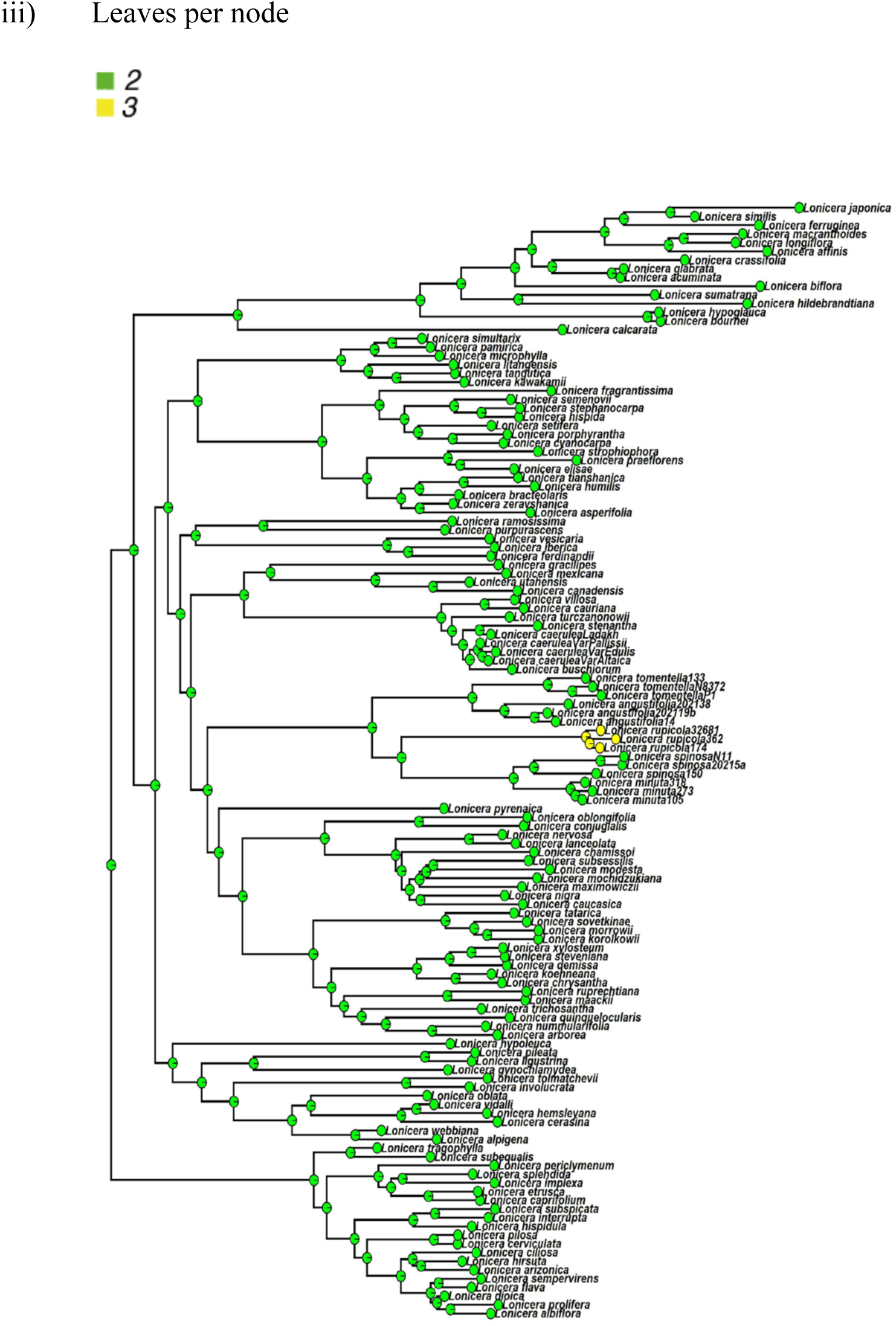
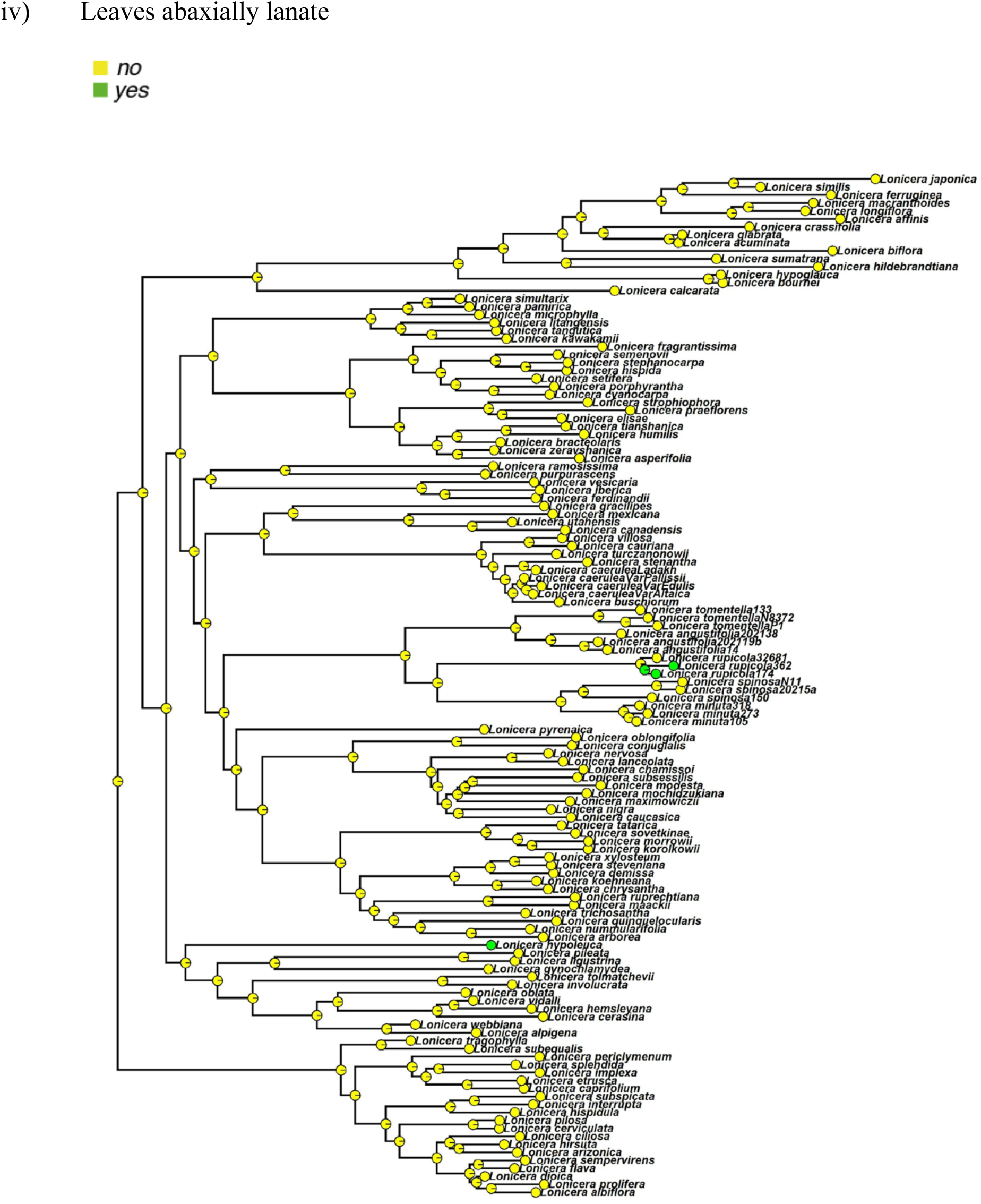
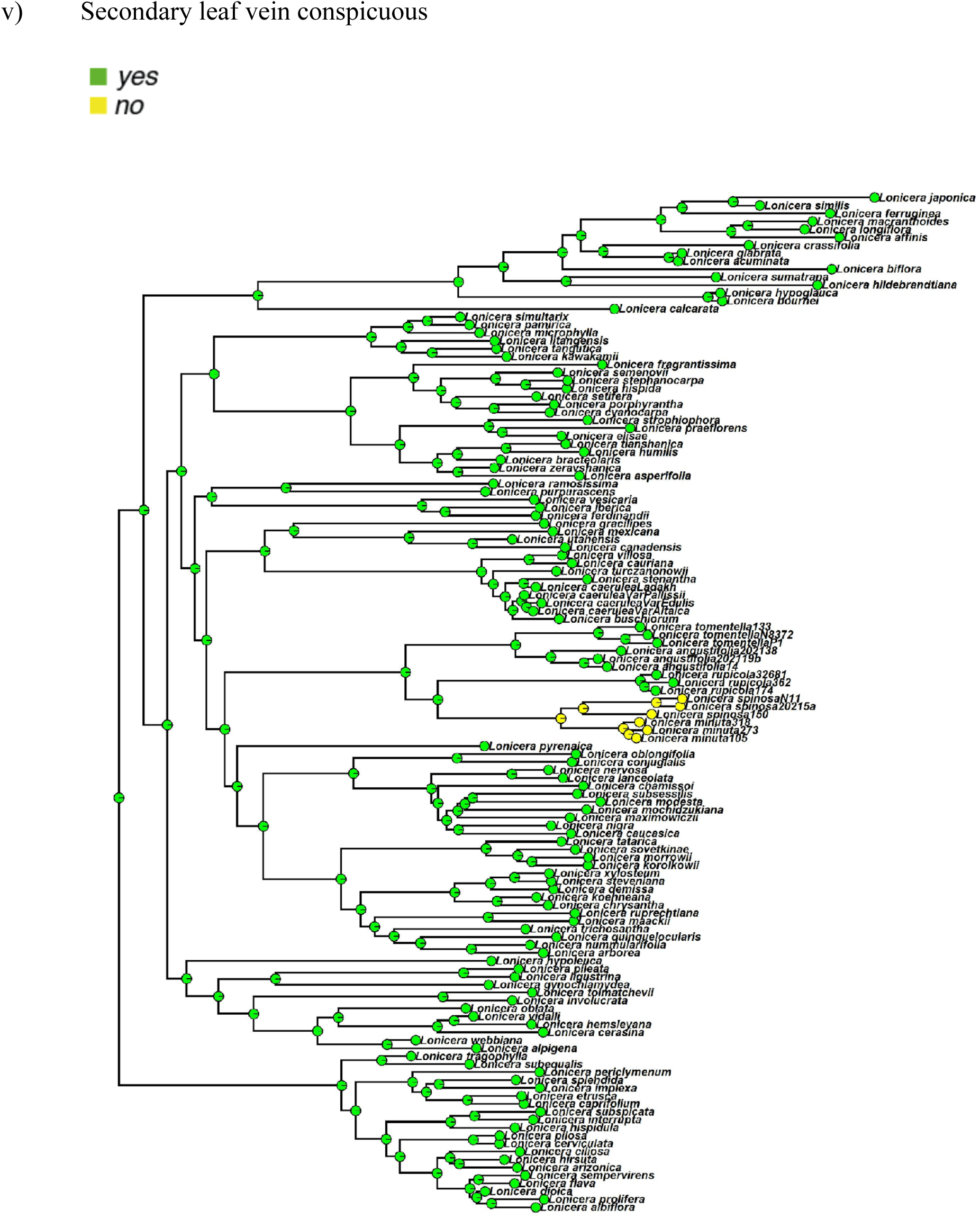
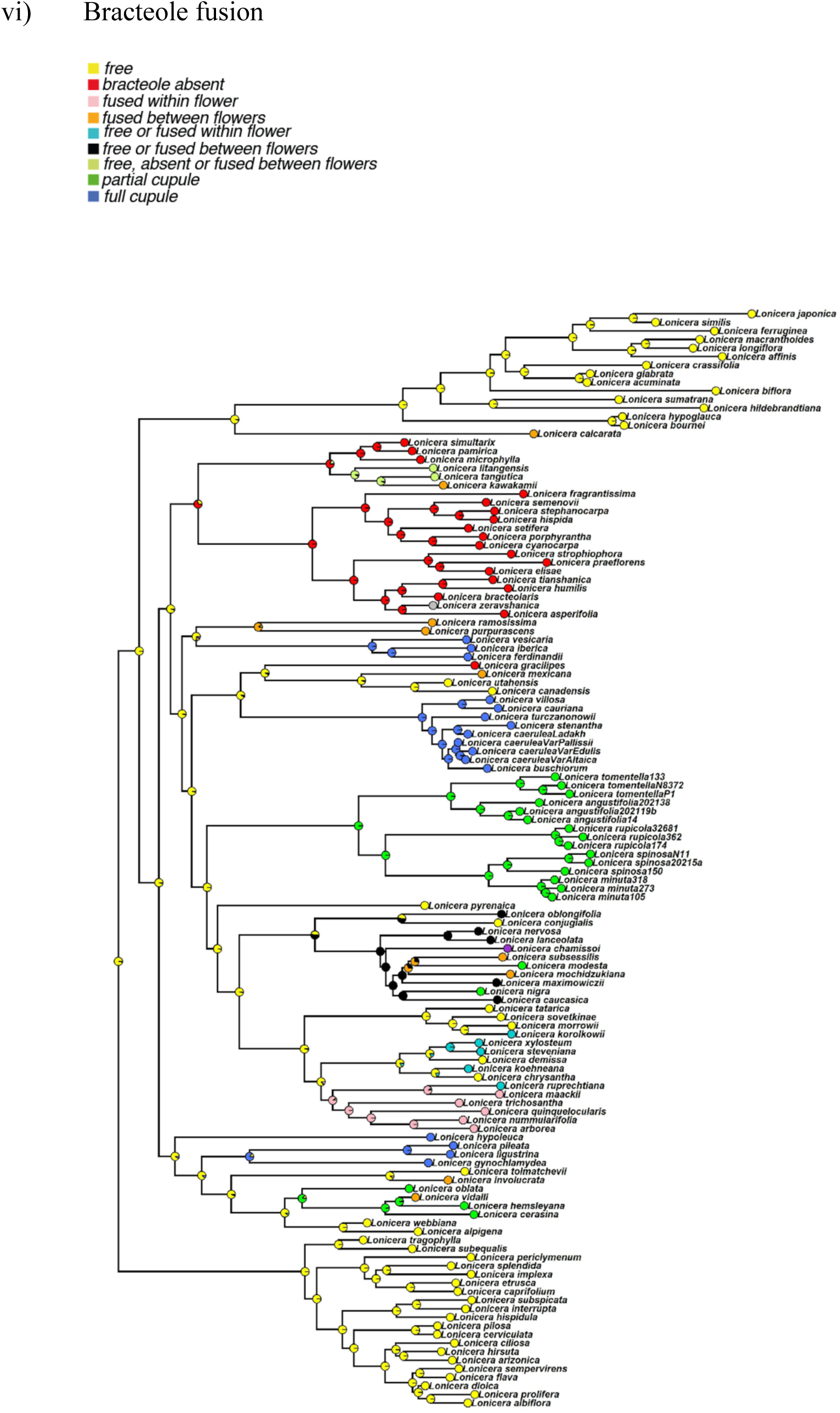
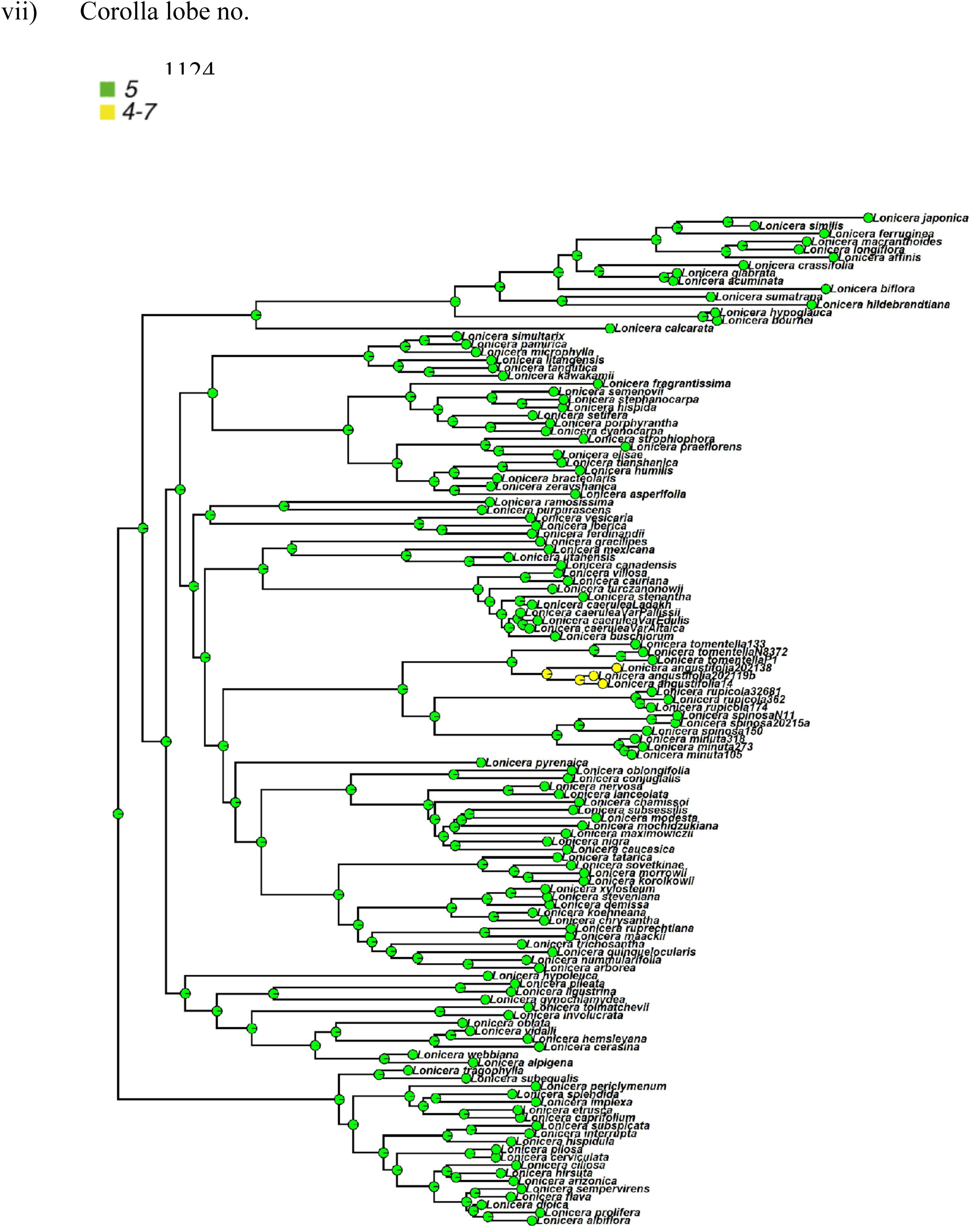
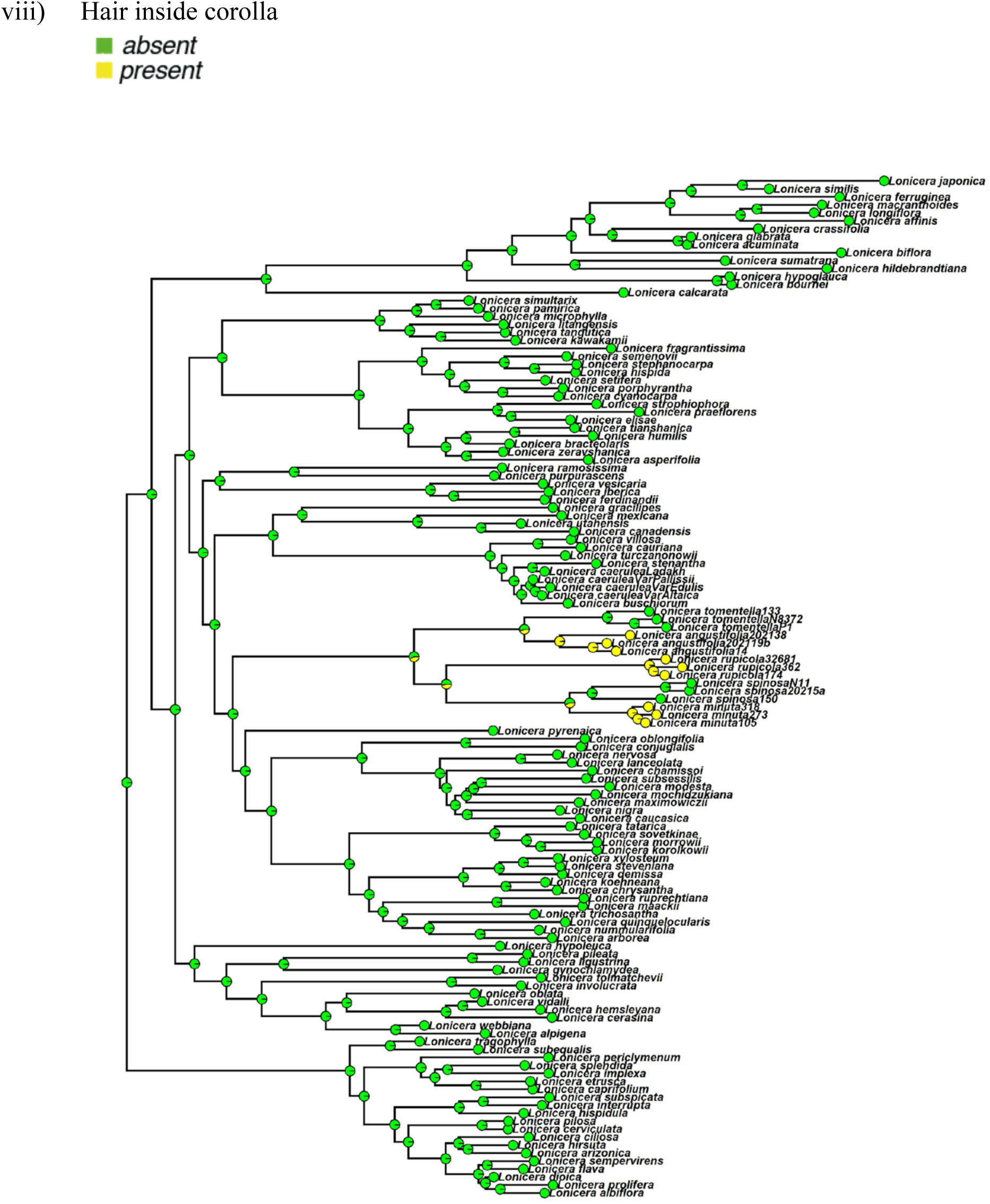
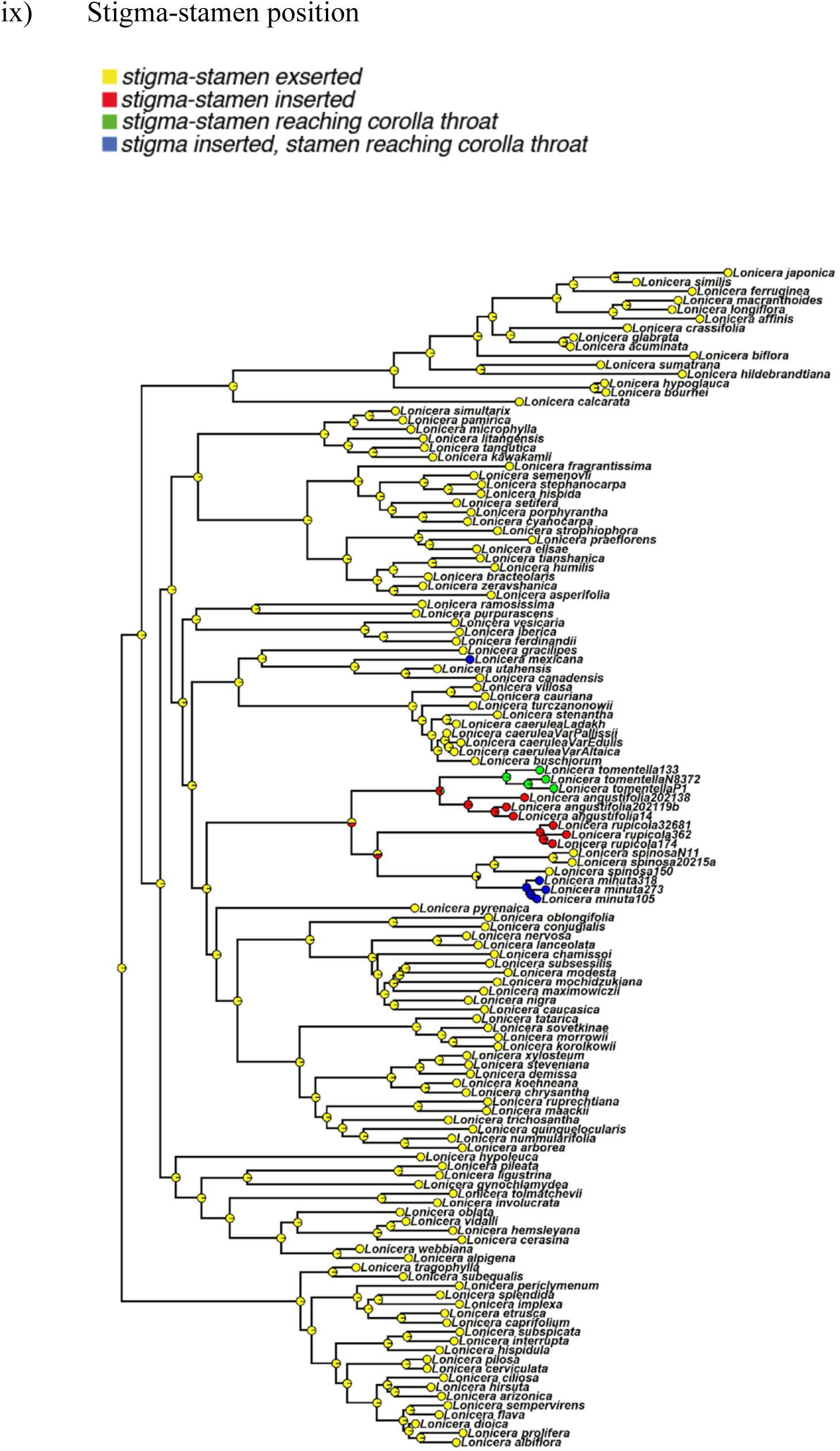
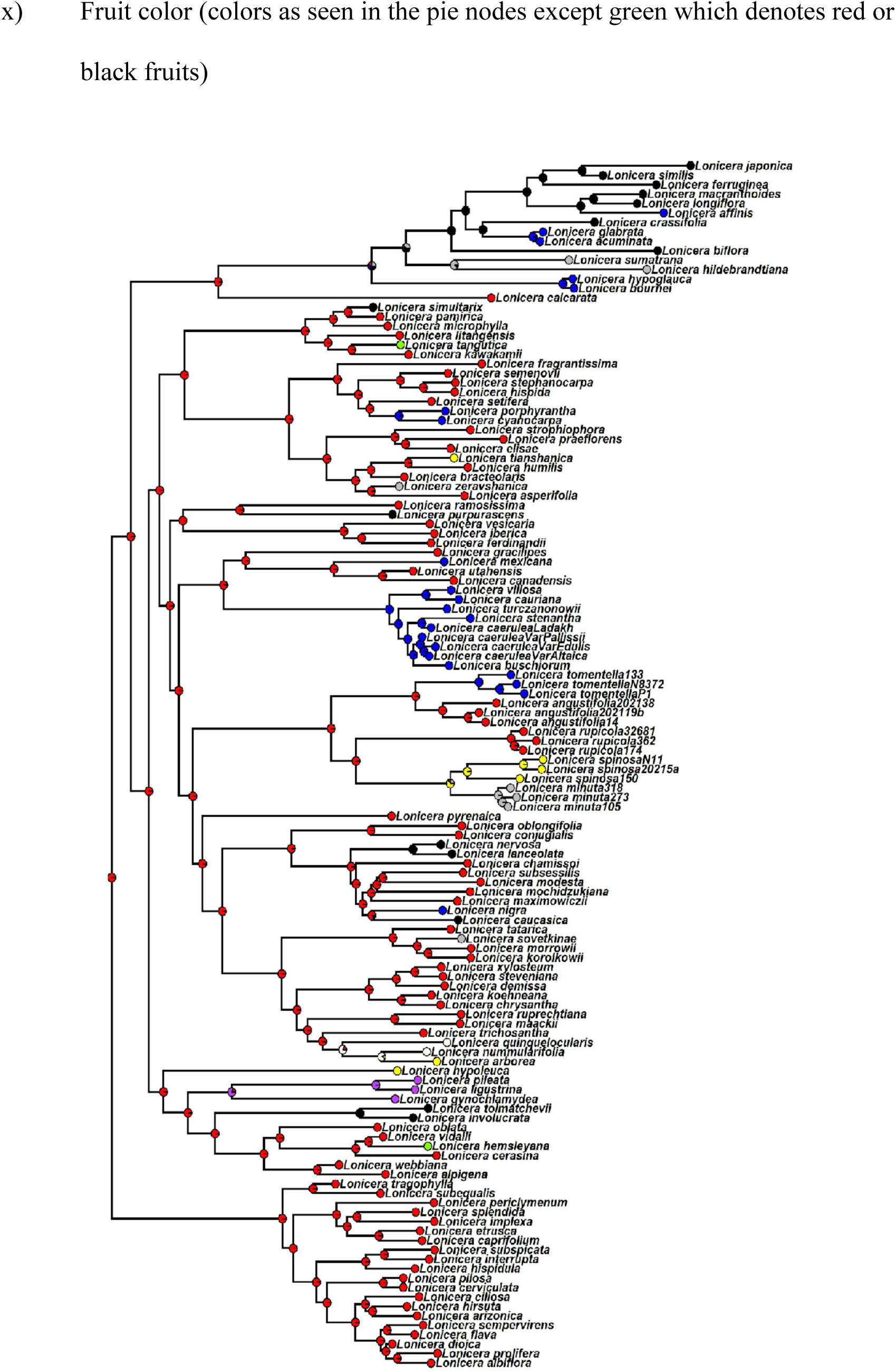
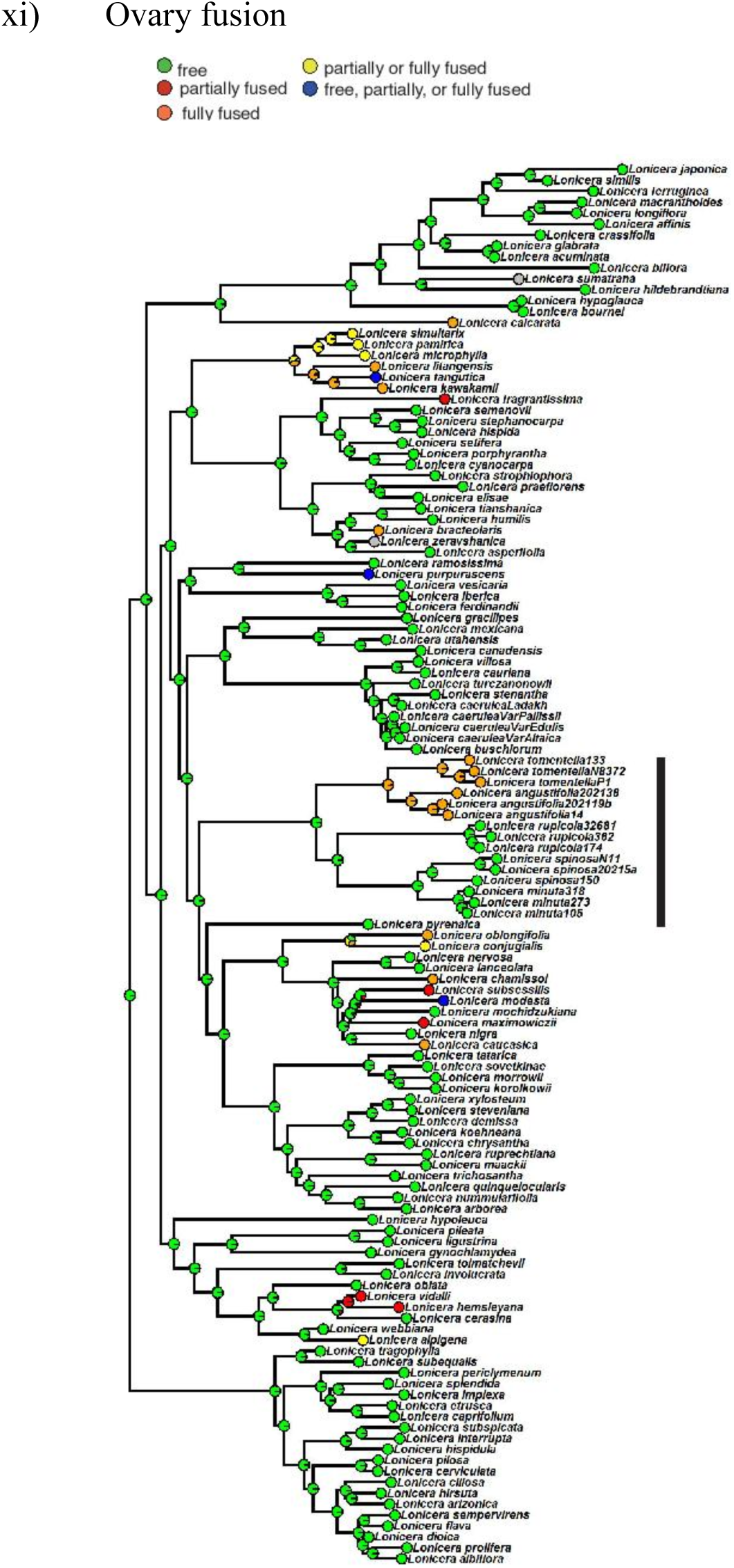
Ancestral state reconstructions of seledcted morphological traits in *Isoxylosteum*.

**FIGURE S7.**
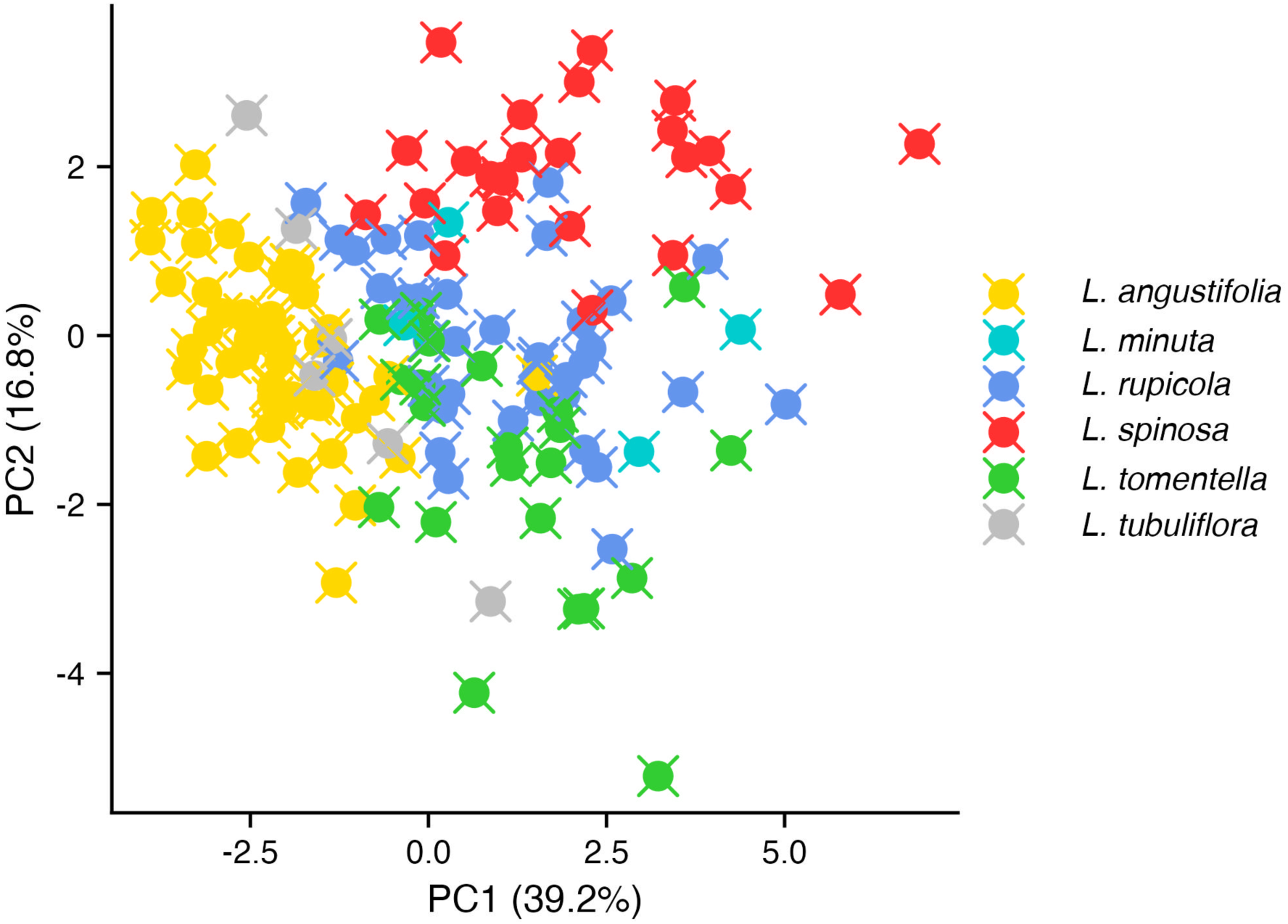
PCA of floral traits with *L. tubuliflora* included.

